# Neurobehavioral effects of fungicides in zebrafish: a systematic review and meta-analysis

**DOI:** 10.1101/2023.06.06.543927

**Authors:** Carlos G. Reis, Leonardo M. Bastos, Rafael Chitolina, Matheus Gallas-Lopes, Querusche K. Zanona, Sofia Z. Becker, Ana P. Herrmann, Angelo Piato

## Abstract

With the aim of yielding high productivity levels, pesticides are widely used in global agriculture. Among them, fungicides are compounds intended to inhibit fungal proliferation in crops and seeds. Their application often leads to environmental contamination, with these chemicals persistently being detected in surface waters. This presence may threaten non-target organisms that dwell in the affected ecosystems, including humans. In toxicologic research, the zebrafish (*Danio rerio*) is the most used fish species to assess the potential effects of fungicide exposure, generating numerous and sometimes conflicting findings. We conducted a systematic review and meta-analysis aiming to synthesize the neurobehavioral effects of fungicides in zebrafish. The search was performed in three databases (PubMed, Scopus, and Web of Science) and the screening was based on a two-stage process guided by pre-defined inclusion/exclusion criteria. Qualitative and quantitative data, as well as reporting quality, were extracted from the included studies (n = 60). Meta-analyses were performed for the outcomes of distance traveled in larvae and adults, and spontaneous movements in embryos. We found an overall significant effect of fungicide exposure on distance, which was lower in exposed versus control groups (SMD −0.44 [−0.74; −0.13], p = 0.0055). No effect was observed for spontaneous movements. The overall heterogeneity for distance and spontaneous movements was considered high (I^2^ = 80%) and moderate (I^2^ = 74%), respectively. This can be explained by substantial methodological variation between protocols, whereas a poor reporting practice hinders the proper critical evaluation of the findings. However, a sensitivity analysis did not indicate any study skewing the meta-analyses. This review demonstrates the need for better-designed and reported experiments in this field.

**Highlights:**

- We systematically reviewed the behavioral effects of fungicides in zebrafish
- Fungicides decrease the distance traveled
- Fungicide exposure has no significant effects on spontaneous movements
- Moderate to high levels of heterogeneity were found
- The results showed a need for better-designed studies with clarity of report

## 1. Introduction

Chemical pesticides are synthetical active ingredients used to control pests that may threaten the productivity of crops (FAO and WHO, 2022). With the aim of yielding high productivity levels, modern agriculture employs large amounts of pesticides (Sharma et al., 2019). In 2020, the global consumption of these products reached almost 3 million tonnes (FAOSTAT, 2023). The substantial quantity and the method by which they are applied results in environmental contamination of the soil, surface waters, and food (Maggi et al., 2019; Pignati et al., 2017; Tudi et al., 2021). Data shows that only less than 0.1% of the pesticide hits the intended target species, leaving the remaining residual as an impact on the environment and public health (Pimentel, 1995). Its presence in superficial waters generates risk to the non-target organisms by decreasing biodiversity and the population of primary producers of the food chain as well as reducing the prey for the aquatic organisms (Beketov et al., 2013; Clasen et al., 2018; McMahon et al., 2012). Moreover, the dissemination of pesticides in the environment represents a risk to humans, whereas their presence in the water supply leads to potential drinking and bioaccumulation in the animals consumed in our diet (Akoto et al., 2016; Albuquerque et al., 2016; de Araújo et al., 2022).

According to the target organism, these substances can be classified as herbicides, insecticides, rodenticides, and fungicides (Delgado-Blanca et al., 2019), being the fungicides one of the most used chemicals (Zubrod et al., 2019). Their application aims to kill and/or inhibit fungal growth in agriculture, both in seeds and crops (Beckerman et al., 2023).

Due to the need to understand the effects of the exposure to these products, the scientific literature presents several studies with animals in this area (J. R. Richardson et al., 2019). The model organism zebrafish (*Danio rerio,* Hamilton 1822) is widely used in toxicology, mostly because of its high fecundity, fast development, transparency of the embryo, and high homology of organs and genetics concerning humans (Bakkers, 2011; Dai et al., 2014; Howe et al., 2013). In addition, the zebrafish is an aquatic animal that dwells in potentially contaminated ecosystems, hence representing the eventual consequences of the exposure to other cohabitant species (Fitzgerald et al., 2021). It has been reported that exposure to fungicides in zebrafish causes behavioral, neurochemical, developmental, metabolic, hormonal, hepatotoxic, cardiotoxic, enzymatic, morphological, and molecular alterations (Cao et al., 2019a; Jia et al., 2020; Jiang et al., 2020; Meng et al., 2020; Shen et al., 2020; Teng et al., 2020; Wang et al., 2020).

From 2012 to 2019, the number of articles that investigate the effects of this type of pesticide on zebrafish occupies the second position in comparison to the others (Gonçalves et al., 2020). However, there is a high methodological heterogeneity between the studies. The interventions, developmental stages, and outcomes addressed are extremely variable between studies. Regarding the intervention, there are plenty of compounds used as fungicides that exhibit distinct mechanisms of action (Lushchak et al., 2018) and can be administered over a wide range of durations through multiple routes of administration. As for the developmental stage, *in vivo* exposure can be performed at different times of the life cycle of the animal (embryo, larval or adult); And the outcomes are distinctly selected according with the research question and the capabilities of the research group (neurotoxicity, hepatotoxicity, cardiotoxicity, among others) (Hamm et al., 2019).

Many studies were published on the toxic effects of fungicides on neurobehavioral parameters in zebrafish (Fitzgerald et al., 2021; Yanicostas and Soussi-Yanicostas, 2021). However, there are no secondary studies that systematically synthesize these results, in order to obtain an understanding supported by published evidence to optimize the planning of new research. An accurate description of these preclinical data, together with a meta-analysis, can help to avoid redundant studies and the consequent use of animals. Furthermore, considering the reproducibility issues also raised for the zebrafish research field (Frommlet, 2020; Gerlai, 2019), it is essential to identify possible sources of bias and conflicting results, which includes assessing the quality of the available publications. This systematic review and meta-analysis of literature aimed at synthesizing the data from neurobehavioral effects of fungicide exposure in zebrafish, also analyzing reporting quality and publication bias.

## 2. Material and methods

Before the screening of studies and data extraction, a protocol guiding this review was registered in Open Science Framework and a pre-registration is available at https://osf.io/y6k5g (Reis et al., 2021). The reporting of this study follows the Preferred Reporting Items for Systematic Reviews and Meta-Analyses (PRISMA) guidelines (Page et al., 2021).

### 2.1) Search strategy

The studies were identified through a search in the literature using three different databases: PubMed, Scopus, and Web of Science. The search strategies were designed in order to adapt to the characteristics of each database. The terms were combined for the intervention (fungicide exposure) and for the population of interest (zebrafish), aiming to conduct a comprehensive search, which included all the available articles that fulfilled the inclusion criteria. The complete query for each database can be found at https://osf.io/5ae9q (Reis et al., 2021). The strategy did not apply any search filter, language restriction, or limit of year. The search was performed on the 1^st^ of December, 2021 and the articles were imported to Rayyan software (Ouzzani et al., 2016) in order to identify and remove the duplicates.

### 2.2) Study screening

Initially, the retrieved studies from the three databases were analyzed to filter and exclude duplicates (CGR). The remaining articles were pre-selected based on their title and abstract. If a reason to exclude the record was not found, at this stage, it was carried forward to the full-text screening stage. In both stages (title/abstract and full-text) two independent reviewers (CGR and LMB, RC or SZB) examined each study. Disagreements between the decisions of the reviewers were resolved by a third reviewer (QKZ, AP, or APH).

Experimental studies evaluating the effects of exposure to fungicides in zebrafish on the following parameters were included: motor function, sensory function, learning and memory, social behavior, sexual behavior, eating behavior, anxiety-like or fear-related behaviors, behaviors related to the reward system, and behaviors related to circadian rhythms. The parameters were included only if they were linked to the central nervous system. The identity of the compound as a fungicide was consulted in the Pesticide Properties Database (AERU, 2022).

In the first phase (screening of title/abstract), papers were excluded according to the following criteria:

1. Type of study: reviews, comments, abstracts published in conference proceedings, corrections, editorials;
2. Type of population: *in vitro* investigations or studies with species other than zebrafish;
3. Type of intervention: biological and commercial formulations or mixtures of fungicides, non-interventional studies; In the next phase (full-text screening) the following criteria were added and the articles were excluded based on the above items plus:
4. Control group: when there is no proper control group (same organism, same procedure, except for fungicide exposure);
5. Outcome measures: if there is no assessment of any previously cited neurobehavioral outcome.

More information about this section is available at https://osf.io/wmsvg.

### 2.3) Data extraction

The data extraction was performed by two independent investigators (CGR and LMB, RC or SZB) and disagreements were resolved by a discussion between the two reviewers. The information and values of interest were directly extracted from the text and tables. When not possible, WebPlotDigitizer software (v4.5, Rohatgi, A., Pacifica, CA, USA, https://automeris.io/WebPlotDigitizer) was used to manually determine the values from the graphs. The following data were extracted: (1) study identification: study title, digital object identifier (DOI), first author, last author, year of publication, and last author affiliation; (2) model animal specifications: strain, sex, developmental stage during exposure, age during exposure, developmental stage during test, age during test; (3) fungicide exposure characteristics: fungicide, administration route and type (i.e. static, semi-static or flow through), frequency of renewal, frequency of exposure, duration of exposure, dose/concentration and the interval between exposure and test; (4) test properties: test nomenclature, category of measured variable (e.g., anxiety, locomotor, social) and the measured variable.

Regarding the authors of the studies, co-authorship networks were elaborated using VOSviewer software version 1.6.18 (https://www.vosviewer.com) (van Eck and Waltman, 2010, 2007).

Data were collected for each variable according to the outcomes of interest, including the mean and the number of animals (n) for both the control and exposed groups. The standard deviation (SD) or standard error of the mean (SEM) was extracted for the reported mean value. If the SEM was reported, the SD was calculated multiplying SEM by the square root of the sample size (SD = SEM ∗ √n).

In instances where the sample size was reported as a range, the lowest value was used. Whenever information was unclear or missing, attempts were made to contact the corresponding author of the study via email, with two separate attempts made at least two weeks apart.

### 2.4) Reporting quality

For the purpose of assessing the reporting quality of included studies, two independent reviewers (CGR and LMB, RC, or SZB) evaluated each paper based on Landis et al., 2012, which proposes criteria for transparent reporting. The observed topics were: (1) mention of any randomization process; (2) sample size estimation; (3) mention of inclusion/exclusion criteria; (4) mention of any process to ensure blinding during the experiments. A score of “yes” or “no” was given for each topic, meaning that it was or was not reported, respectively. The outcome measurements performed by any automated software were considered blinded. Reporting quality plots were created using robvis (McGuinness and Higgins, 2021).

A complete guide for assessing the reporting quality associated with each of the items in this review is available at https://osf.io/uy5v3.

### 2.5) Meta-analysis

To perform a meta-analysis, a requirement of at least 5 studies with the same outcome was defined a priori (Reis et al., 2021). Whenever two or more experimental groups shared the same control, the sample size of the control group was divided by the number of comparisons and then rounded down (Vesterinen et al., 2014).

Effects sizes were determined with standardized mean differences (SMD) using Hedge’s G method. Analyses were conducted using R Project for Statistical Computing with packages meta (Balduzzi et al., 2019) (https://cran.r-project.org/package=meta) and ggplot2 (Wilkinson, 2011) following Hedge’s random effects model given the anticipated heterogeneity between studies. Values for SMD were reported with 95% confidence intervals. Heterogeneity between studies was estimated using I² (Higgins and Thompson, 2002), τ², and Cochran’s Q (Cochran, 1954) tests. Heterogeneity variance (τ²) was estimated using the restricted maximum likelihood estimator (Veroniki et al., 2016; Viechtbauer, 2005). The confidence intervals around pooled effects were corrected using Knapp-Hartung adjustments (Knapp and Hartung, 2003). Values of 25%, 50%, and 75% were considered as representing low, moderate, and high heterogeneity, respectively for I², and a p-value ≤ 0.1 was considered significant for Cochran’s Q. Prediction intervals were estimated and represent the range of effects expected for future studies (Higgins et al., 2019). Furthermore, a subgroup meta-analysis was performed to evaluate if the developmental stage of the animals was a potential source of heterogeneity. Studies were grouped into two categories: larval and adult. Subgroup analysis was only performed when there were at least five unique studies for each subgroup. A p-value ≤ 0.1 was considered significant for subgroup differences (M. Richardson et al., 2019).

To investigate an association between the effect and the class of fungicide, we conducted an exploratory meta-analysis by categorizing the fungicides based on their chemical structure. Even without reaching the minimum of 5 studies, we ran a meta-analysis with 4 articles that investigated fungicides of the triazole and anilide groups.

To explore the relationship between the effect sizes and fungicide concentration as a moderator variable, a mixed-effects meta-regression analysis was conducted. The random effects structure accounted for potential heterogeneity across studies (Viechtbauer et al., 2015). Meta-regressions excluding studies based on the sensitivity analysis were also performed.

Publication bias was investigated by generating funnel plots and performing Duval and Tweedie’s trim and fill analysis (Duval and Tweedie, 2000) and Egger’s regression test (Egger et al., 1997). Analyses were only conducted when at least five studies were available within a given outcome for funnel plots and at least ten studies for the regression test. A p-value < 0.1 was considered significant for the regression test.

#### 2.5.1) Sensitivity analysis

A sensitivity analysis was conducted to assess if any experimental or methodological difference between studies was biasing the main effect found in the meta-analysis. Analyses were performed following the “leave-one-out jackknife method” (Miller, 1974). A minimum of three comparisons were required for each outcome in order to conduct a sensitivity analysis. Furthermore, we conducted complementary meta-analyzes excluding studies that, when omitted in the leave-one-out, had observations that changed the overall effect direction. We also ran meta-analyses excluding studies containing experiments with atypically high SMD as seen in the forest plots (Andrade, 2020).

## 3. Results

### 3.1) Search results

The search in the three databases retrieved a total of 2139 results. After the removal of duplicates, 1140 articles were screened for eligibility by analyzing the titles and abstracts. As a result of the first screening phase, 369 studies remained to be assessed based on their full-text. At this phase, 3 were not retrieved and 60 fulfilled the criteria and were included in the review (Fig. 1). The main overall reasons for the exclusions were outcome (n = 234), population (n = 260) and intervention (n = 350). Concerning the quantitative synthesis, a total of 8 studies were excluded because the minimum number of studies to perform a meta-analysis was not reached for the reported outcomes and 10 because of missing information. There were 18 experiments measuring distance using luminous transitions (dark/light) in larvae that were not included due to the variations between the protocols (Hill et al., 2023), which makes the comparison unfeasible. This resulted in 24 studies included in the quantitative synthesis. Detailed reasons for the exclusion of studies from the meta-analysis are available at https://osf.io/qpcew.

**Fig. 1.**
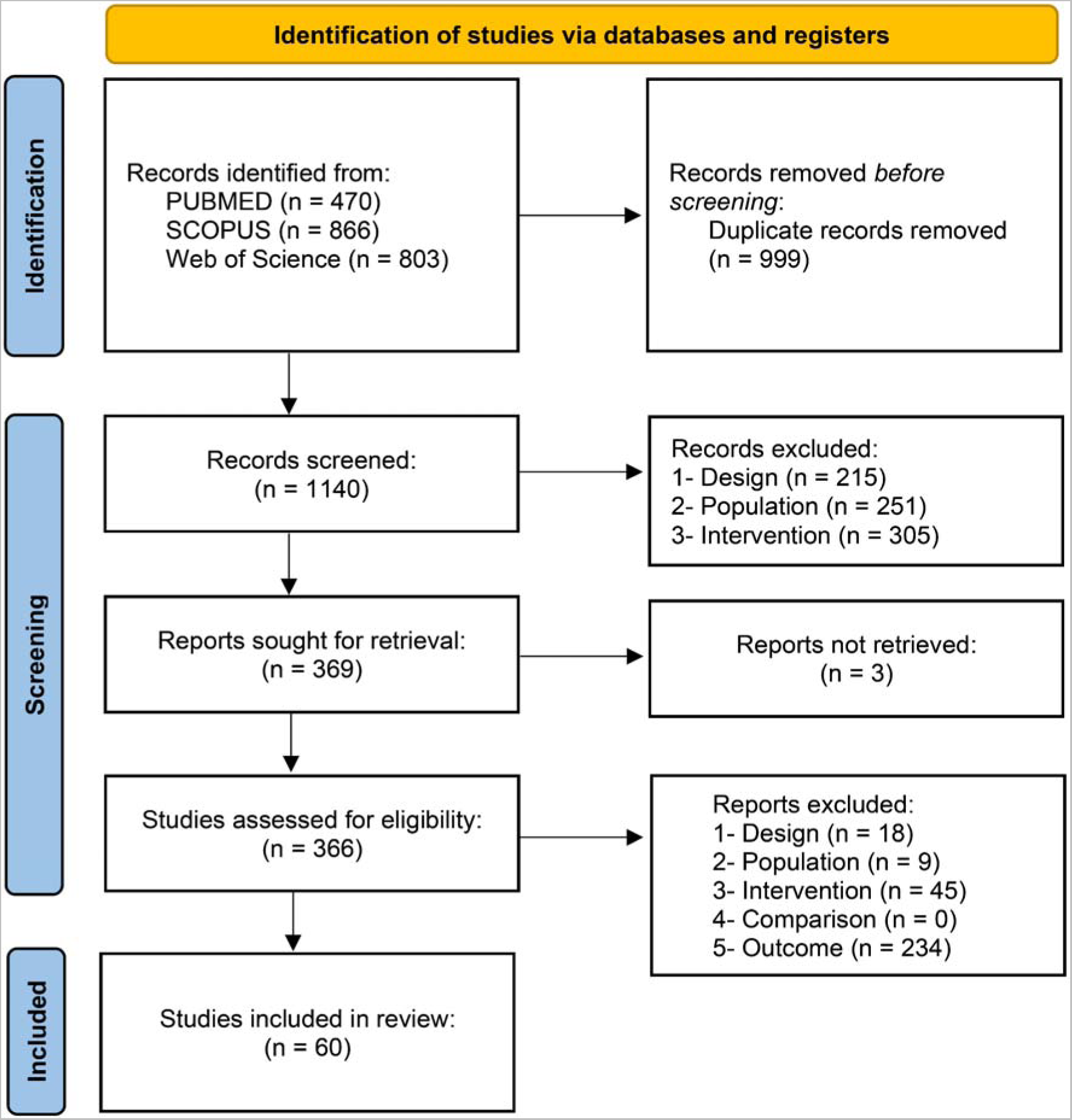
Flowchart diagram of the collection of studies and selection process.

### 3.2) Study characteristics

A qualitative description of the studies is provided in Table 1. A total of 43 different fungicides were addressed in the articles included in this review. Studies with the fungicides difenoconazole (n = 5, 8.3%), boscalid (n = 4, 6.6%) and pyraclostrobin (n = 4, 6.6%) were the most frequent.

**Table 1.** Qualitative description of studies reporting effects of the exposure to fungicides on neurobehavioral and neurochemical outcomes in zebrafish. The indicated concentrations of exposure were used to assess the behavioral outcomes. The main findings were described as: ↑, higher when compared to the control group; ↓, lower when compared to the control group; =, no difference when compared to the control group.

| Study ID | Fungicide | Concentration (mg/L) | Duration of exposure | Developmental stage during exposure/outcome assessment | Main findings |
| --- | --- | --- | --- | --- | --- |
| Domingues, 2013 | Prochloraz | 0.3, 0.6, 1.2, 2.4, 4.8, 7.2, 9.6 | 96 h | Embryo/Larvae & Adult/Adult | **Locomotor behavior**<br>↑ Spontaneous movements<br>↓ Distance traveled<br>↑ Distance in the dark<br>↑ Distance in the light<br>↑ Velocity<br>↑ Acceleration<br>↑ Absolut turn angle<br>**Neurochemical outcomes**<br>↓ ChE activity (larvae)<br>↑ GST activity (larvae) |
| Fitzmaurice, 2013 | Benomyl | 0.29 | 120 h | Embryo/Larvae | **Locomotor behavior**<br>↓ Distance traveled |
| Mu, 2013 | Difenoconazole | 0.5, 1, 1.5, 2, 2.5, 3 | 24 h | Embryo/Embryo | **Locomotor behavior**<br>↑ Spontaneous movements (1.5 mg/L)<br>↓ Spontaneous movements (2.5, 3 mg/L)<br>↑ Reversal rate behavior |
| Andrade, 2016 | Carbendazim | 1.1, 1.19, 1.3, 1.41, 1.53, 1.66, 1.8 | 120 h | Embryo/Larvae | **Locomotor behavior**<br>↓ Distance in the light<br>↓ % Small distance in the light<br>↓ % Small distance in the dark<br>↑ % Long distance in the light<br>↑ % Long distance in the dark<br>↑ Swimming time in the light<br>↑ Swimming time in the dark<br>**Neurochemical outcomes**<br>↑ ChE activity<br>↑ GST activity<br>= CAT activity |
| Jin, 2016 | Imazalil | 0.01, 0.03, 0.1, 0.3 | 96 h | Embryo/Larvae | **Locomotor behavior**<br>↓ Distance traveled<br>↓ Distance in the dark<br>↓ Distance in the light<br>↓ Velocity<br>**Neurochemical outcomes**<br>↓ AChE levels<br>↓ AChE activity<br>= DA levels |
| Lulla, 2016 | Ziram | 0.0003 - 0.305 | 7 days | Embryo/Larvae | **Locomotor behavior**<br>↓ Distance in the dark<br>= Distance in the light<br>↓ Velocity |
| Mu, 2016 | Difenoconazole | 0.5, 2 | 96 h | Embryo/Embryo | **Locomotor behavior**<br>= Spontaneous movements |
| Yang, 2016 | Thifluzamide | 2.66, 2.76, 2.85, 2.95,<br>3.04, 3.23<br>&<br>2.66, 2.76, 2.85, 2.95,<br>3.04<br>&<br>2.66, 2.85, 3.04, 3.23,<br>3.42, 3.61 | 96 h<br>&<br>144 h<br>&<br>96 h | Embryo/Embryo<br>&<br>Embryo/Larvae<br>&<br>Larvae/Larvae | **Locomotor behavior**<br>↓ Spontaneous movements (embryo)<br>↓ Swimming rate (larvae) |
| Yang, 2016b | Flutolanil | 1.5, 1.8, 2.16, 2.59, 3.1 | 24 h | Embryo/Embryo | **Locomotor behavior**<br>↑ Spontaneous movements |
| Altenhofen, 2017 | Tebuconazole | 1, 2, 4<br>&<br>1, 4, 6 | 120 h<br>&<br>96 h | Embryo/Larvae<br>&<br>Adult/Adult | **Locomotor behavior**<br>↓ Distance traveled<br>↓ Absolut turn angle (larvae)<br>= Crossings<br>**Anxiety/fear related behavior**<br>↓ Time in the periphery<br>= Time in the upper zone<br>**Aggressive behavior**<br>↑ Time in the bottom<br>**Neurochemical outcomes**<br>↓ AChE activity |
| De la Paz, 2017 | Triadimefon | 16 | 8 h | Larvae/Larvae | **Locomotor behavior**<br>↑ Locomotor activity |
| Costa-Silva, 2018 | Mancozeb | 1 | 23 h<br>&<br>43 h | Embryo/Embryo<br>&<br>Embryo/Embryo<br>dechorionated | **Locomotor behavior**<br>↑ Spontaneous movements<br>↑ Number of stimulus (embryo<br>dechorionated)<br>↑ Response to touch (embryo dechorionated)<br>**Neurochemical outcomes**<br>= GST activity<br>↓ GSH levels<br>= GPx activity |
| Fan, 2018 | Hymexazol | 417, 480, 554, 639, 738 | 48 h | Embryo/Larvae | **Locomotor behavior**<br>↓ Swimming rate<br>**Neurochemical outcomes**<br>↑ % Inhibition of tyrosinase activity |
| Li, 2018 | Pyraoxystrobin | 2.03, 2.44, 2.9, 3.51, 4.22, 5.08 | 24 h | Embryo/Embryo | **Locomotor behavior**<br>= Spontaneous movements |
| Qian, 2018 | Boscalid | 0.7, 2, 2.3, 2.6, 2.9, 3.2 | 22 h | Embryo/Embryo | **Locomotor behavior**<br>↑ Spontaneous movements |
| Teng, 2018 | Difenoconazole | 0.0005, 0.005, 0.05, 0.5 | 24 h | Embryo/Embryo | **Locomotor behavior**<br>↑ Spontaneous movements |
| Teng, 2018b | Difenoconazole | 0.0005, 0.005, 0.05, 0.5 | 24 h | Embryo/Embryo | **Locomotor behavior**<br>↑ Spontaneous movements |
| Wang, 2018 | Fluazinam | 0.04, 0.09, 0.13 | 6 days | Embryo/Larvae | **Locomotor behavior**<br>↑ Swimming activity in the dark (0.04 mg/L)<br>↓ Swimming activity in the dark (0.09, 0.13 mg/L)<br>= Swimming activity in the light |
| Cao, 2019 | Ziram | 0.0003, 0.003 | 7 days | Embryo/Larvae | **Locomotor behavior**<br>↑ Swimming activity<br>↑ Distance traveled (0.003 mg/L)<br>= Distance in the dark<br>= Distance in the light<br>= Total velocity<br>↑ Velocity in light<br>**Anxiety/fear related behavior**<br>↓ Time in the dark<br>= Frequency in the dark |
| Cao, 2019b | Cyproconazole | 2.9, 7.2, 14.5, 29.1, 72.9, 145.8 | 24 h<br>&<br>48 h<br>&<br>7 days | Embryo/Embryo<br>&<br>Embryo/Larvae | **Locomotor behavior**<br>↓ Spontaneous movements<br>↓ Swimming activity in the dark<br>= Swimming activity in the light |
| Cao, 2019c | Maneb | 0.02, 0.13, 0.26 | 7 days | Embryo/Larvae | **Locomotor behavior**<br>↓ Swimming activity in the dark<br>↓ Swimming activity in the light |
| Li, 2019 | Pyraclostrobin | 0.009, 0.018, 0.36 | 4 days | Larvae/Larvae | **Locomotor behavior**<br>↓ Distance traveled<br>↓ Velocity<br>**Neurochemical outcomes**<br>↑ Glutamate receptor activity |
| Paredes-Zúñiga, 2019 | Triadimefon | 5, 20, 35 | 10 h<br>&<br>11 min | Larvae/Larvae<br>&<br>Adult/Adult | **Locomotor behavior**<br>↓ Swimming activity<br>↓ Distance traveled (larvae)<br>↑ Distance traveled (adult)<br>↑ Velocity<br>**Anxiety/fear related behavior**<br>↓ Time in the periphery<br>↑ Time in the bottom zone<br>↓ Time in the upper zone<br>**Aggressive behavior**<br>↑ Number of bites<br>**Neurochemical outcomes**<br>↑ DA levels<br>↓ 5-HT levels |
| Perez-Rodriguez, 2019 | Tebuconazole | 0.03, 0.3, 3 | 6 days | Embryo/Larvae | **Locomotor behavior**<br>↓ Distance in the dark<br>= Distance in the light<br>= Velocity<br>**Anxiety/fear related behavior**<br>↑ Mean time in the dark<br>= Cumulative time in the dark<br>↑ Frequency in the dark zone |
| Qian, 2019 | Penthiopyrad | 2.3, 2.4, 2.5, 2.6, 2.7, 2.8, 2.9<br>&<br>0.3, 0.6, 1.2 | 1 day<br>&<br>5-8 days | Embryo/Embryo<br>&<br>Embryo/Larvae | **Locomotor behavior**<br>↑ Spontaneous movements (2.5, 2.6, 2.7 mg/L)<br>↓ Spontaneous movements (2.9 mg/L)<br>↓ Swimming activity<br>↓ Distance traveled<br>↓ Velocity<br>↓ Acceleration |
| Souders, 2019 | Propiconazole | 0.03, 0.3, 3.4 | 144 h | Embryo/Larvae | **Locomotor behavior**<br>↓ Distance traveled<br>↓ Distance in the dark |
| Teng, 2019 | Propiconazole | 0.5, 2.5, 4.5 | 24 h<br>&<br>120 h | Embryo/Embryo<br>&<br>Embryo/Larvae | **Locomotor behavior**<br>↑ Spontaneous movements<br>↓ Distance traveled<br>↓ Velocity<br>↓ Swimming activity<br>↓ Acceleration |
| Tian, 2019 | Prothioconazole | 0.0375, 0.075, 0.15 | 24 h | Embryo/Embryo | **Locomotor behavior**<br>= Spontaneous movements<br>**Neurochemical outcomes**<br>↓ GSH levels<br>= SOD activity<br>= CAT activity<br>↑ MDA levels |
| Valadas, 2019 | Propiconazole | 0.000425, 0.00085,<br>0.0017, 0.0085 | 96 h | Adult/Adult | **Locomotor behavior**<br>= Distance traveled<br>↓ Crossings<br>**Anxiety/fear related behavior**<br>↓ Time in the upper zone<br>↑ Time in the upper zone<br>↓ Entries in the upper zone<br>= Entries in the bottom zone<br>**Neurochemical outcomes**<br>↑ SOD activity<br>↑ CAT activity<br>= MDA levels<br>= SH levels<br>= NPSH levels |
| Wang, 2019 | Oxine-copper | 0.01, 0.02, 0.04 | 24 h | Embryo/Embryo<br>&<br>Larvae/Larvae | **Locomotor behavior**<br>↓ Number of tail coiling<br>↓ Distance traveled<br>↓ Swimming activity<br>↓ Velocity<br>↑ Absolut turn angle<br>**Neurochemical outcomes**<br>↓ AChE activity(embryo)<br>↑ SOD activity (embryo)<br>↑ CAT activity (embryo)<br>↑ MDA levels (embryo)<br>↑ ROS levels (embryo) |
| Yang, 2019 | Flutolanil | 0.125, 0.5, 2 | 24 h<br>&<br>96 h | Embryo/Embryo<br>&<br>Embryo/Larvae | **Locomotor behavior**<br>↓ Spontaneous movements<br>= Distance traveled<br>**Neurochemical outcomes**<br>↑ DA levels |
| Yang, 2019b | Thiﬂuzamide | 0.19, 1.9, 2.85 | 24 h<br>&<br>96 h | Embryo/Embryo<br>&<br>Embryo/Larvae | **Locomotor behavior**<br>= Spontaneous movements<br>= Distance traveled<br>**Neurochemical outcomes**<br>↓ DA levels |
| Zhou, 2019 | Captan | 0.58, 0.66, 0.75, 0.86,<br>1.00, 1.16 | 24 h | Embryo/Embryo | **Locomotor behavior**<br>= Spontaneous movements |
| Hussain, 2020 | Tebuconazole<br>&<br>Dimethomorph<br>&<br>Difenoconazole | 0.3<br>&<br>0.3<br>&<br>0.4 | 24 h | Larvae/Larvae | **Locomotor behavior**<br>↑ Distance in the dark (tebuconazole,<br>dimethomorph)<br>↓ Distance in the dark (difenoconazole)<br>↑ Distance in the light<br>↑ Burst movement count in the light<br>↑ Burst movement count in the dark<br>(dimethomorph, difenoconazole)<br>↑ Rotation count in the light (dimethomorph,<br>difenoconazole)<br>↑ Rotation count in the dark (dimethomorph,<br>difenoconazole) |
| Jia, 2020 | Penconazole (+)<br>&<br>Penconazole (-) | 1, 2 | 24 h<br>&<br>96 h | Embryo/Embryo<br>&<br>Embryo/Larvae | **Locomotor behavior**<br>↑ Spontaneous movements ((+)-<br>penconazole)<br>↓ Velocity ((+)-penconazole)<br>**Neurochemical outcomes**<br>↓ AChE activity ((+)-penconazole)<br>↓ DA levels ((+)-penconazole)<br>↓ 5-HT levels ((+)-penconazole)<br>= Glycine levels<br>= Norepinephrine levels |
| Kumar, 2020 | Azoxystrobin<br>&<br>Pyraclostrobin | 0.00001, 0.0001, 0.01,<br>0.1, 1 | 5 days | Embryo/Larvae | **Locomotor behavior**<br>↓ Distance traveled<br>**Neurochemical outcomes**<br>= MDA levels |
| Liu, 2020 | Propamocarb | 0.01, 0.1, 1 | 7 days | Embryo/Larvae | **Locomotor behavior**<br>↑ Distance traveled<br>↑ Distance in the dark<br>= Distance in the light<br>↑ Velocity<br>**Neurochemical outcomes**<br>↓ AChE activity<br>= MDA levels<br>↓ DA levels<br>= SOD activity<br>↑ CAT activity<br>↑ GPx activity<br>↓ GST activity |
| Pang, 2020 | Myclobutanil | 4, 6, 8, 10, 12, 14, 16 | 24 h | Embryo/Embryo | **Locomotor behavior**<br>↑ Spontaneous movements (4, 6, 8, 10,<br>12 mg/L)<br>↓ Spontaneous movements (16 mg/L) |
| Shen, 2020 | Mepanipyrim | 0.0001, 0.001, 0.01, 0.1 | 7 days | Embryo/Larvae | **Locomotor behavior**<br>↑ Distance traveled (7, 14 dpf)<br>↓ Distance traveled (14 dpf)<br>↑ Velocity (7, 14 dpf)<br>↓ Velocity (14 dpf)<br>↑ Acceleration<br>= Absolut turn angle<br>↓ Immobile time (7 dpf)<br>**Neurochemical outcomes**<br>= AChE activity<br>↑ GABA levels |
| Souders, 2020 | Triticonazole | 0.3, 3.1, 31.7 | 6 days | Embryo/Larvae | **Locomotor behavior**<br>↑ Distance in the dark<br>= Distance in the light |
| Tang, 2020 | Cyprodinil | 0.0001, 0.001, 0.01, 0.1 | 24 h | Embryo/Embryo | **Locomotor behavior**<br>↓ Spontaneous movements |
| Teng, 2020 | Flutolanil | 0.00025, 0.05, 1 | 60 days | Adult/Embryo (offspring) | **Locomotor behavior**<br>= Spontaneous movements |
| Vasamsetti, 2020 | Etridiazole | 3.75, 7.5, 15, 30, 60 | 96 h | Embryo/Larvae | **Locomotor behavior**<br>↑ Immobile time |
| Wang, 2020 | Boscalid | 5, 15, 25 | 24 h | Embryo/Embryo<br>&<br>Larvae/Larvae | **Locomotor behavior**<br>↓ Number of tail coiling<br>↓ Distance traveled<br>↓ Distance in the dark<br>↓ Distance in the light<br>↓ Velocity<br>↑ Absolut turn angle<br>↑ Immobile time<br>**Neurochemical outcomes**<br>= AChE activity<br>↑ MDA levels<br>↓ SOD activity<br>↑ CAT activity<br>↑ ROS levels |
| Zhang, 2020 | Zoxamide | 0.16, 0.33, 0.84, 1.68 | 24 h<br>&<br>6 days | Embryo/Larvae<br>&<br>Embryo/Larvae | **Locomotor behavior**<br>↑ Distance in the dark (24 h, 6 days exposure)<br>↓ Distance in the dark (6 days exposure)<br>↑ Distance in the light (6 days exposure) |
| Barreto, 2021 | Fosetyl-al | 0.02, 0.2, 2, 20, 200 | 120 h | Embryo/Larvae | **Locomotor behavior**<br>↓ Distance traveled<br>= Swimming time<br>↑ Velocity<br>↑ Acceleration<br>↑ Absolut turn angle<br>**Neurochemical outcomes**<br>= ChE activity<br>= CAT activity<br>↑ GST activity |
| Brenet, 2021 | Bixafen | 0.08, 0.2 | 96 h | Embryo/Larvae | **Locomotor behavior**<br>↓ Distance traveled |
| Fan, 2021 | Carbendazim | 0.52, 0.65, 0.82, 1.02, 1.28, 1.6 | 24 h | Embryo/Embryo | **Locomotor behavior**<br>↑ Spontaneous movements |
| Forner-Piquer, 2021 | Boscalid<br>&<br>Captan<br>&<br>Thiophanate<br>&<br>Ziram | 0.00001, 0.00005, 0.001, 0.01, 0.1, 1, 10 | 120 h | Embryo/Larvae | **Locomotor behavior**<br>↓ Distance traveled<br>↓ Velocity |
| Huang, 2021 | Fenamidone | 0.03, 0.3, 0.4, 0.6 | 144 h | Embryo/Larvae | **Locomotor behavior**<br>= Distance traveled (light-dark test)<br>↓ Distance in the dark (visual motor response test)<br>= Distance in the light<br>**Anxiety/fear related behavior**<br>= Frequency in the dark<br>= Time in the dark |
| Leandro, 2021 | Mancozeb | 0.005, 0.01, 0.02 | 24 h<br>&<br>68 h<br>&<br>164 h | Embryo/Embryo<br>&<br>Embryo/Larvae | **Locomotor behavior**<br>↓ Spontaneous movements (28 hpf)<br>↑ Number of stimulus (72 hpf)<br>↓ Response to touch (72 hpf)<br>↑ Distance traveled (0.005 mg/L, 168 hpf)<br>↓ Distance traveled (0.02 mg/L, 168 hpf)<br>↑ Absolut turn angle (0.005 mg/L, 168 hpf)<br>↓ Absolut turn angle (0.02 mg/L, 168 hpf)<br>↑ Immobile episodes (0.02 mg/L, 168 hpf)<br>↓ Immobile episodes (0.005 mg/L, 168 hpf)<br>↑ Immobile time (0.02 mg/L, 168 hpf)<br>↓ Immobile time (0.005 mg/L, 168 hpf)<br>**Anxiety/fear related behavior**<br>= Time in the periphery (168 hpf)<br>↑ Entries in the periphery (168 hpf)<br>**Neurochemical outcomes**<br>↑ AChE activity (28 hpf)<br>↓ AChE activity (72 hpf)<br>↓ SOD activity (24 hpf)<br>↑ SOD activity (72 hpf)<br>↓ CAT activity (72 hpf)<br>↑ CAT activity (168 hpf)<br>↑ GST activity (72 hpf)<br>↓ GST activity (168 hpf)<br>↑ ROS levels (72, 168 hpf) |
| Li, 2021 | Azoxystrobin<br>&<br>Kresoxim-methyl<br>&<br>Pyraclostrobin<br>&<br>Trifloxystrobin | 0.02027<br>&<br>0.01567<br>&<br>0.01939<br>&<br>0.02042 | 5 days | Embryo/Larvae | **Locomotor behavior**<br>↑ Distance in the dark (kresoxim-methyl, pyraclostrobin, trifloxystrobin)<br>↑ Distance in the light (trifloxystrobin)<br>**Neurochemical outcomes**<br>↑ MDA levels (pyraclostrobin, trifloxystrobin)<br>↑ SOD activity (pyraclostrobin, trifloxystrobin)<br>↑ CAT activity (trifloxystrobin)<br>↑ ROS levels (pyraclostrobin, trifloxystrobin) |
| Lin, 2021 | Fluxapyroxad | 1.1, 1.2, 1.3, 1.4, 1.5, 1.6 | 24 h | Embryo/Embryo | **Locomotor behavior**<br>↑ Spontaneous movements<br>**Neurochemical outcomes**<br>↑ MDA levels<br>= SOD activity<br>= CAT activity<br>↑ GPx activity (0.174 mg/L)<br>↓ GPx activity (0.694 mg/L) |
| Paredes-Zúñiga, 2021 | Triadimefon | 5, 15 | 3 days | Adult/Adult | **Locomotor behavior**<br>↑ Time in the drug-paired zone (5 mg/L)<br>↓ Time in the drug-paired zone (15 mg/L)<br>↑ Circling behavior (days 1, 2) |
| Pompermaier, 2021 | Copper | 0.105 | 48 h | Adult/Adult | **Locomotor behavior**<br>= Distance traveled<br>= Absolut turn angle<br>= Crossings<br>**Anxiety/fear related behavior**<br>= Time in the upper zone<br>= Time in the middle zone<br>= Time in the bottom zone |
| Qian, 2021 | Boscalid | 0.3, 0.6, 1.2<br>&<br>0.01, 0.1, 1.0 | 8 days<br>&<br>21 days | Embryo/Larvae<br>&<br>Adult/Adult | **Locomotor behavior**<br>↓ Distance traveled (larvae)<br>↑ Distance traveled (adult)<br>↓ Distance in the dark (larvae)<br>↓ Distance in the light (larvae)<br>↓ Velocity<br>↓ Acceleration<br>↓ Active time (larvae)<br>↑ Active time (larvae, adult)<br>**Neurochemical outcomes**<br>↑ ACh levels (larvae)<br>↓ AChE activity (larvae) |
| Tang, 2021 | Cyprodinil | 0.0001, 0.001, 0.01 | 209-211 days<br>&<br>215-217 days | Embryo/Adult | **Locomotor behavior**<br>↓ Distance traveled<br>↓ Velocity<br>= Acceleration<br>↓ Absolut turn angle<br>**Aggressive behavior**<br>↑ Time in the interaction zone |
| Wu, 2021 | Procymidone | 0.001, 0.01, 0.1 | 4 days<br>&<br>7 days | Embryo/Larvae | **Locomotor behavior**<br>↑ Distance in the dark (4 days)<br>↑ Distance in the light (4 days)<br>↓ Distance in the dark (7 days)<br>↓ Distance in the light (7 days) |
| Yang, 2021 | Azoxystrobin<br>&<br>Pyraclostrobin<br>&<br>Trifloxystrobin | 0.0002, 0.001, 0.005<br>&<br>0.77, 1.54, 2.32<br>&<br>0.51, 1, 2 | 6 days | Embryo/Larvae | **Locomotor behavior**<br>↑ Distance in the dark (azoxystrobin, pyraclostrobin)<br>↓ Distance in the dark (trifloxystrobin)<br>= Distance in the light |
| Yang, 2021b | Thiﬂuzamide | 0.19, 1.9, 2.85 | 96 h<br>&<br>144 h | Embryo/Larvae<br>&<br>Embryo/Larvae | **Locomotor behavior**<br>↓ Distance traveled<br>↓ Velocity<br>↓ Swimming activity<br>↓ Rotating frequency (144 hpf)<br>**Neurochemical outcomes**<br>↓ AChE activity<br>↑ 5-HT levels<br>↑ Norepinephrine levels |

All the studies used immersion as the method of exposure, whereas exposure durations ranged from 11 minutes to 217 days. The most recurrent duration of exposure among the publications was 24 hours (n = 21, 35%), followed by 96 hours (n = 12, 20%). It is important to emphasize that 24 h is usually employed to verify the outcome of spontaneous movements, while 96 h is recommended by the Organisation for Economic Co-operation and Development (OECD) to assess acute fish toxicity in protocols 203 (adults) and 236 (embryos) (OECD, 2019, 2013). Regarding the developmental stage during the exposure, the embryonic was the most common (n = 52, 86.7%). Subsequently, the larval stage was observed in 7 studies (11.6%), and the adult stage in 8 studies (13.3%). Some articles used more than one stage for the exposure.

The outcome assessment was mostly performed in larvae (n = 41, 68.33%) and embryos (n = 25, 41.7%). There were studies that assessed the outcomes in more than one developmental stage.

The sex of the adult animals was mainly reported as an equal proportion of male and female (F:M), with the exception of one in which it was not reported (unclear).

Regarding the authors included in this review, co-authorship network analysis identified 24 clusters of researchers investigating the neurobehavioral effects of fungicides across the globe (Fig. 2). An interactive version of the co-authorship network is available at https://tinyurl.com/239thp6t.

**Fig. 2.**
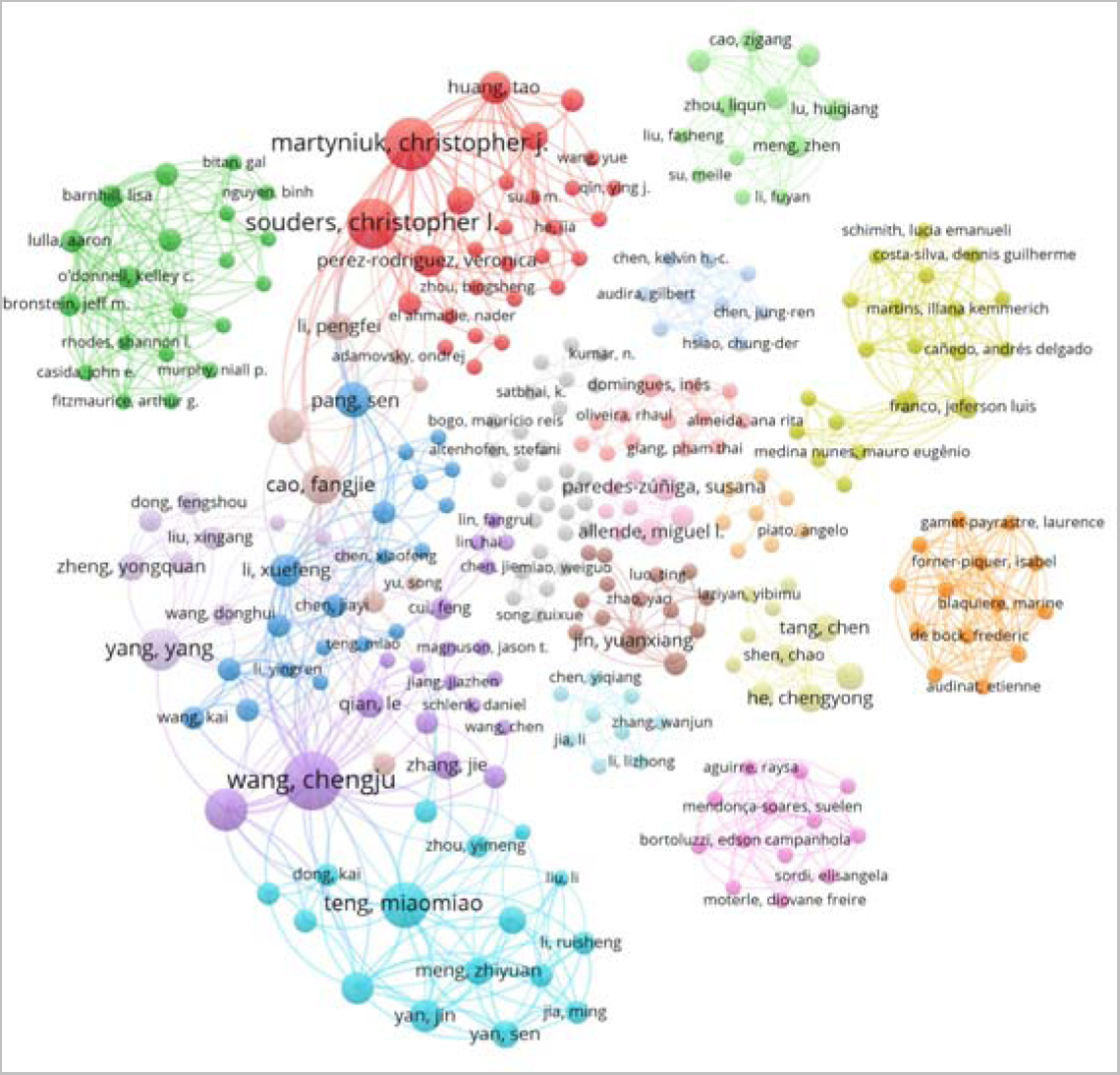
Co-authorship network analysis of researchers who authored the included articles investigating neurobehavioral effects of fungicides in zebrafish. The size of the circles represents the number of studies published by each author. The color of the lines and circles differentiates the clusters of authors. The distance between the two circles indicates the correlations between researchers.

### 3.3) Reporting quality

The summary plot of the reporting quality evaluation is shown in Fig. 3. Randomization process was not cited in 17 studies (28.33%). Only 3 articles described methods for sample size estimation (5%), and none of the authors explicitly stated the data inclusion or exclusion criteria. Blinding was reported in 38 papers (63.33%). Individualized scores for each study included are available at https://osf.io/pgrhq.

**Fig. 3.**
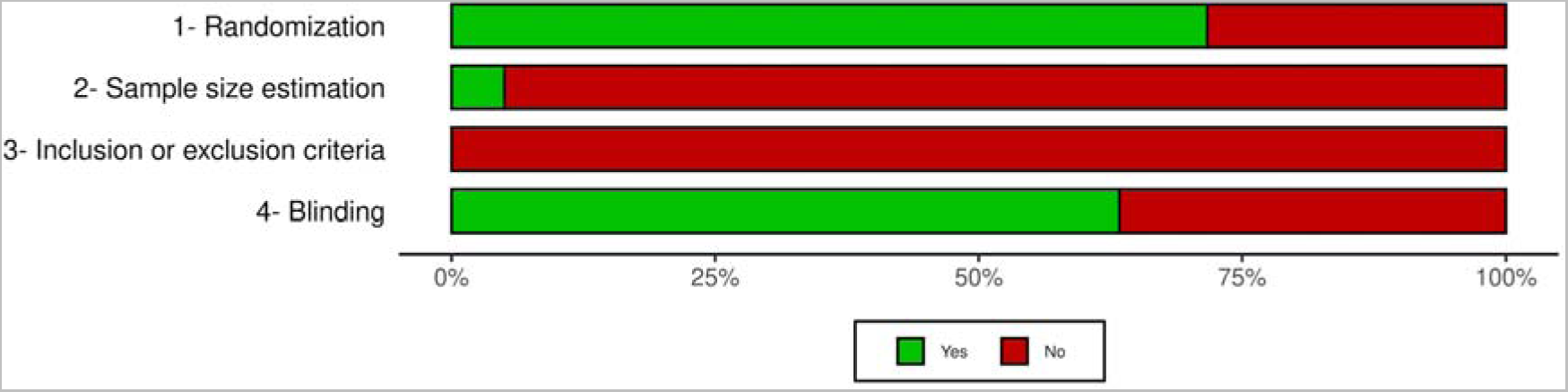
Reporting quality assessment of the included studies. The reporting quality assessment was performed by two independent investigators based on the criteria described by Landis et al., 2012. Each item was scored as yes or no, meaning that the item is either reported or not, respectively. Classification is given as the percentage of assessed studies (n = 60) presenting each score.

### 3.4) Meta-analysis

#### 3.4.1) Distance

The meta-analysis included 61 comparisons from 12 independent studies. The total of animals used as controls was 1112, whereas the exposure individuals counted 2045. For adults, the tests used to assess locomotor behavior were the novel tank test (2), locomotor activity (1), locomotor behavior (1), mirror-induced aggressive behavior (1), and antipredator behavior (1). In larvae, the authors executed locomotor activity (9), locomotor behavior (2), and exploratory behavior (1).

The highest concentration of fungicide in the meta-analysis was 145.89 mg/L for cyproconazole (Cao et al., 2019b), while the lowest was 0.0001 mg/L for mepanipyrim and cyprodinil (Shen et al., 2020; Tang et al., 2020).

The overall analysis showed that exposed animals present a lower distance traveled as compared to controls (SMD −0.44 [−0.74; −0.13], p = 0.0055, Fig. 4). The estimated heterogeneity was considered high, with an I^2^ = 80%, a τ^2^ = 0.88, and a Q = 300.1 (*df* = 60, p < 0.01). When calculating strictly for the developmental stage of the larvae, there was a significant effect of the fungicides on decreasing the distance traveled (SMD −0.44 [−0.83; −0.05], p = 0.03, Fig. 4). The heterogeneity was still considered high for this subgroup, with an I^2^ = 84%, a τ^2^ = 1.21, and a Q = 284.48 (p < 0.01). Similarly, analyzing the adults subgroup there was a significant effect of the exposure to fungicides on decreasing the distance traveled (SMD −0.55 [−0.89; −0.21], p < 0.01, Fig. 4). Unlike the larvae, the heterogeneity was considered low, with an I^2^ = 5%, a τ^2^ = 0.07, and a Q = 13.72 (p = 0.39). The difference between subgroups was not significant (p = 0.68), indicating that the developmental stage is not a direct moderator for this outcome.

**Fig. 4.**
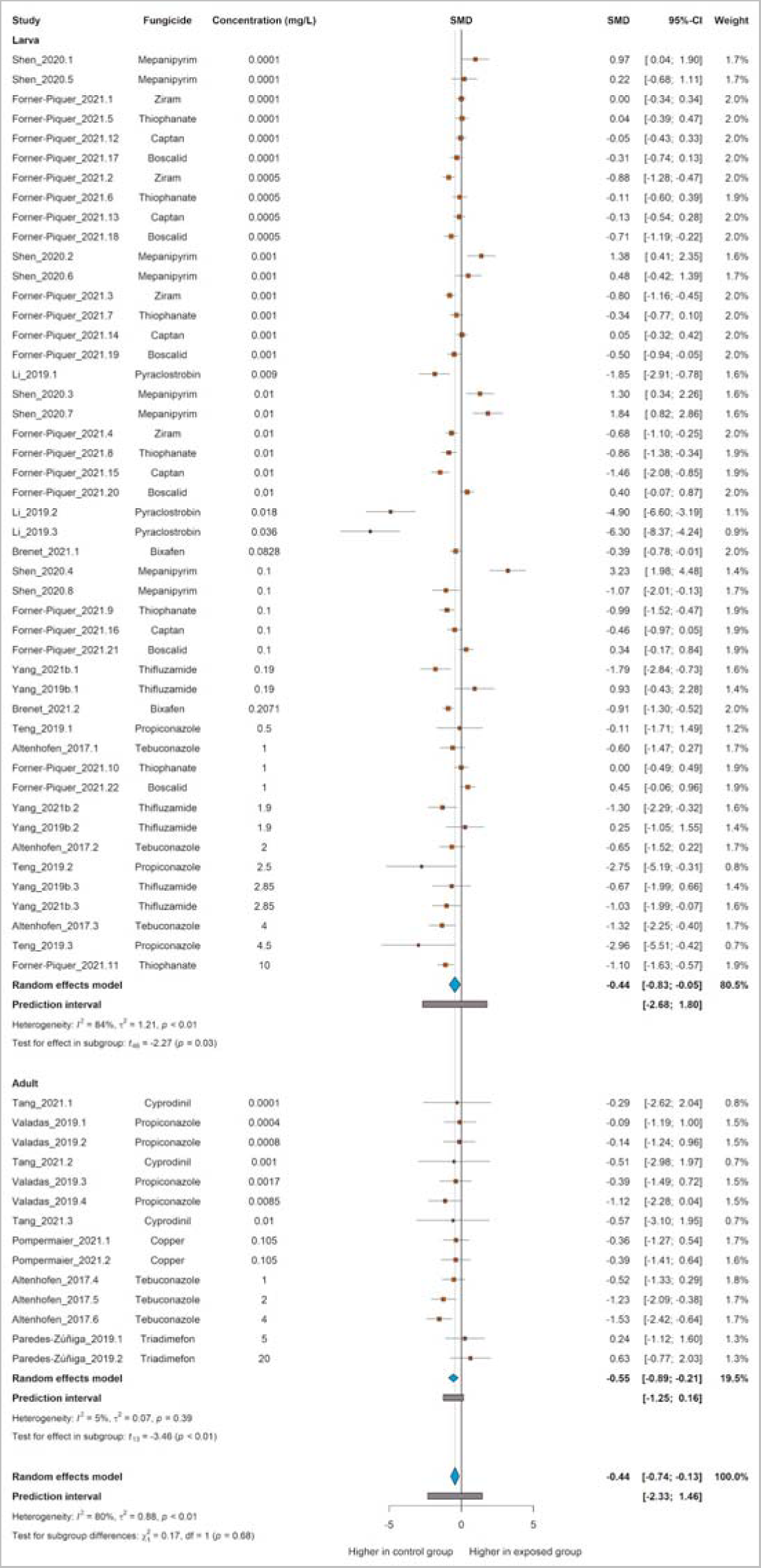
The effect of exposure to fungicides on distance traveled in zebrafish. Subgroup analyses were based on the developmental stage (either larva or adult). Data are presented as Hedges’ G standardized mean differences (SMD) and 95% confidence intervals.

The result from the meta-analysis of distance using only fungicides of the triazole group was similar and a decrease in distance was observed (Fig. S1).

#### 3.4.2) Spontaneous movements

The meta-analysis comprised 64 comparisons from 13 independent studies. The total of embryos used as controls was 190 and the exposure individuals counted 670. All of the experiments performed the outcome assessment at 24 h of exposure, with the exception of one (48 h).

The overall analysis showed that fungicide exposure had no significant effect on the number of spontaneous movements (SMD −0.16 [−0.67; 0.34], p = 0.5265, Fig. 5). The estimated heterogeneity was considered moderate, with an I^2^ = 74%, a τ^2^ = 1.86, and a Q = 243.19 (*df* = 63, p < 0.01).

**Fig. 5.**
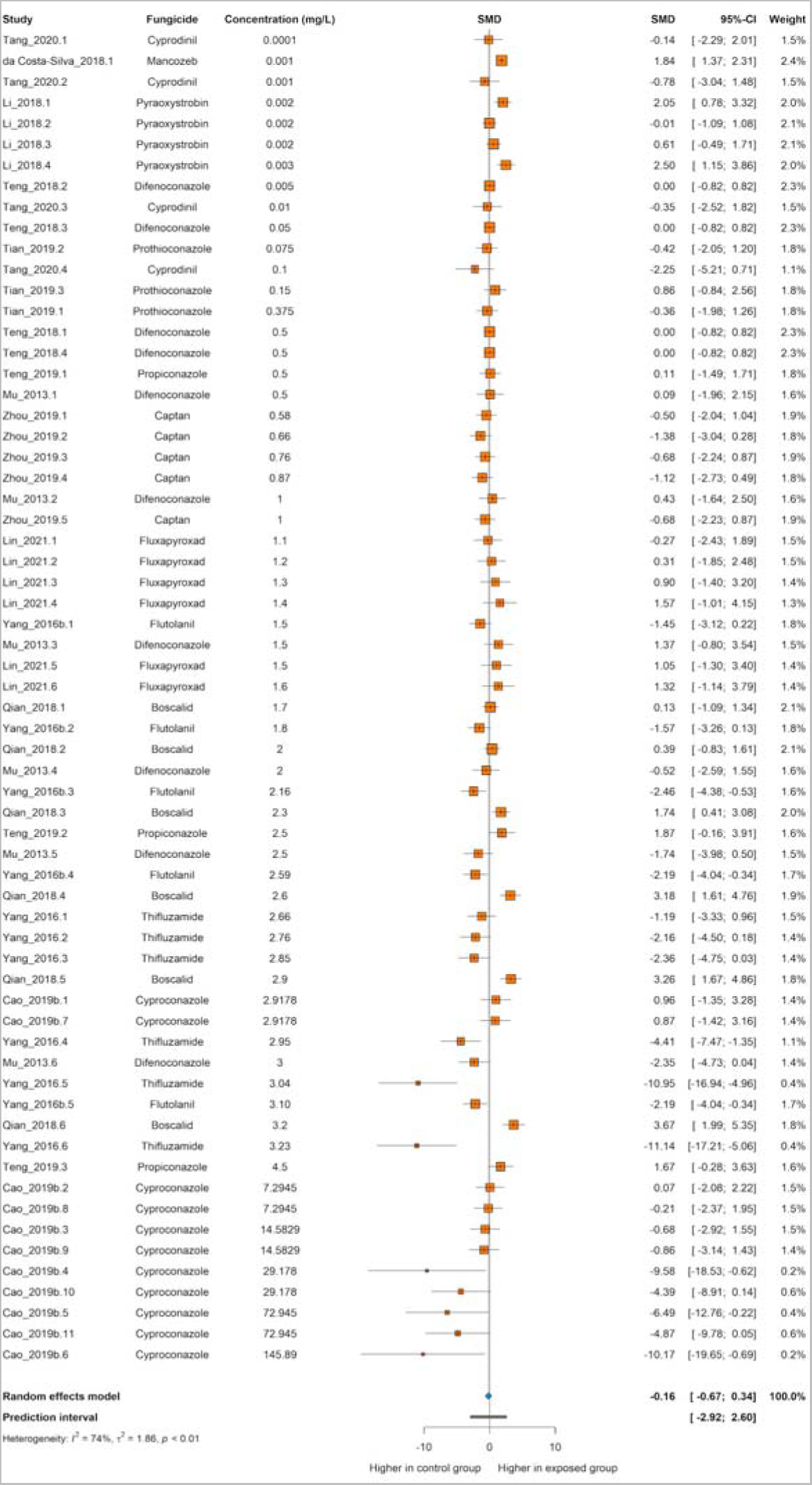
The effect of exposure to fungicides on spontaneous movements in zebrafish. Data are presented as Hedges’ G standardized mean differences (SMD) and 95% confidence intervals.

The result from the meta-analysis of spontaneous movements using only fungicides of the anilide or triazole groups was similar and no significant effects were observed (Fig. S2 and S3, respectively).

The meta-regression of both outcomes showed no significant correlation of the concentration with the effects (Fig. S4 and S5). Meta-regressions excluding studies from Li, 2019 (distance), Cao 2019b and Yang, 2016 (spontaneous movements), maintained no significant correlation (Fig. S6 and S7).

Additional information regarding the meta-analysis can be found at https://osf.io/hdu5c/.

### 3.5) Publication bias

Visual inspection of the funnel plot for the outcome of distance showed the most symmetrical distribution of the studies (Fig. 6) compared to the outcome of spontaneous movements. Trim and fill analysis for distance imputed 4 studies to the meta-analysis, and the overall effect of the fungicides exposure was not significant for this outcome (SMD −0.29 [−0.66, 0.08], p = 0.1252, Fig. 6).

**Fig. 6.**
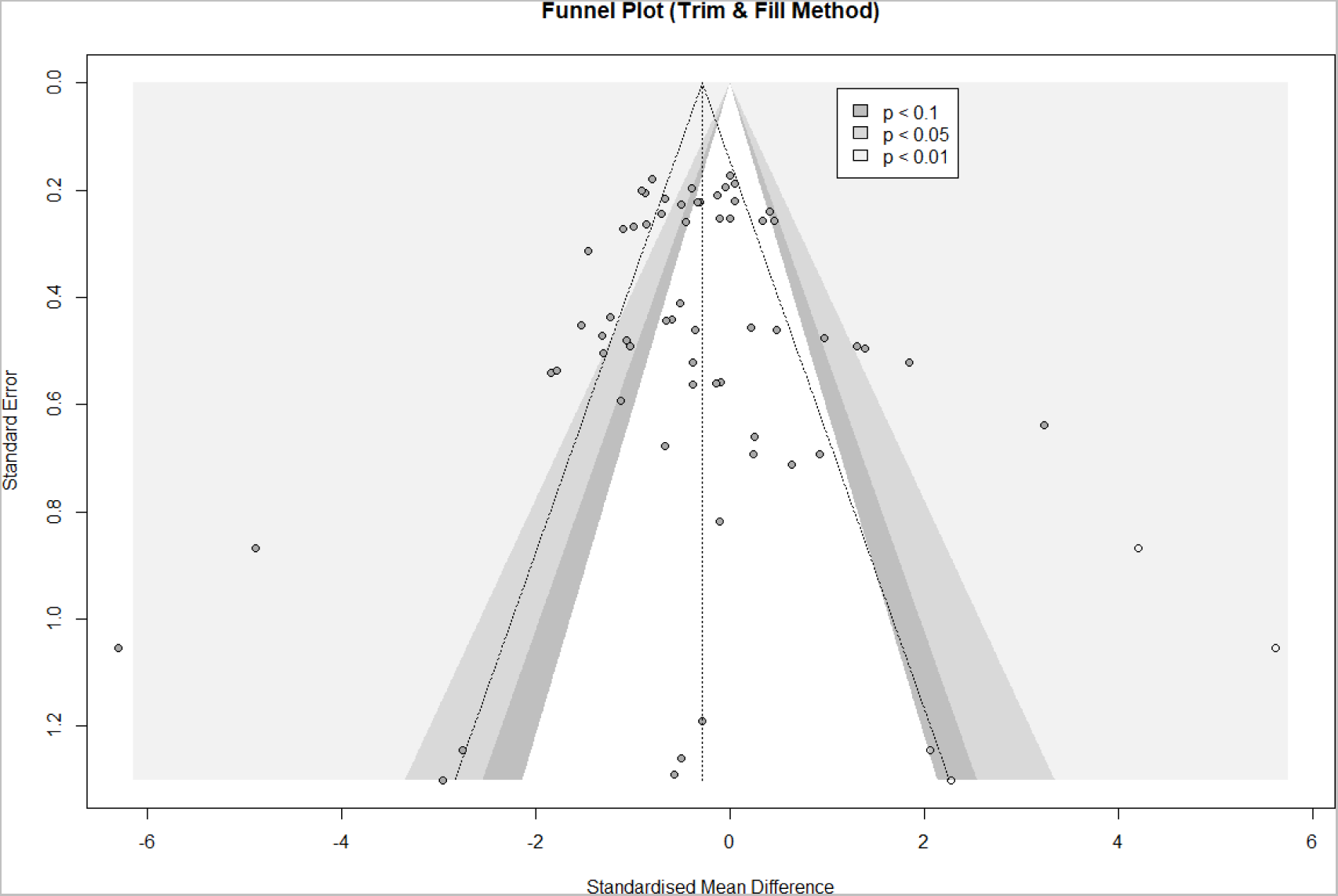
Funnel plot including studies analyzed within the outcome of distance. Each gray circle represents a single comparison. Hollow circles represent imputed studies in the trim and fill analysis. The vertical line represents the overall effect size and the triangular region represents the 95% confidence interval. Shaded areas represent the interval for statistically significant effects.

For spontaneous movements, the funnel plot demonstrated an asymmetrical distribution (Fig. 7). Trim and fill analysis for this outcome imputed 20 studies to the meta-analysis, and the overall effect of the fungicides exposure was not significant (SMD 0.64 [−0.02, 1.29], p = 0.0568, Fig. 7).

**Fig. 7.**
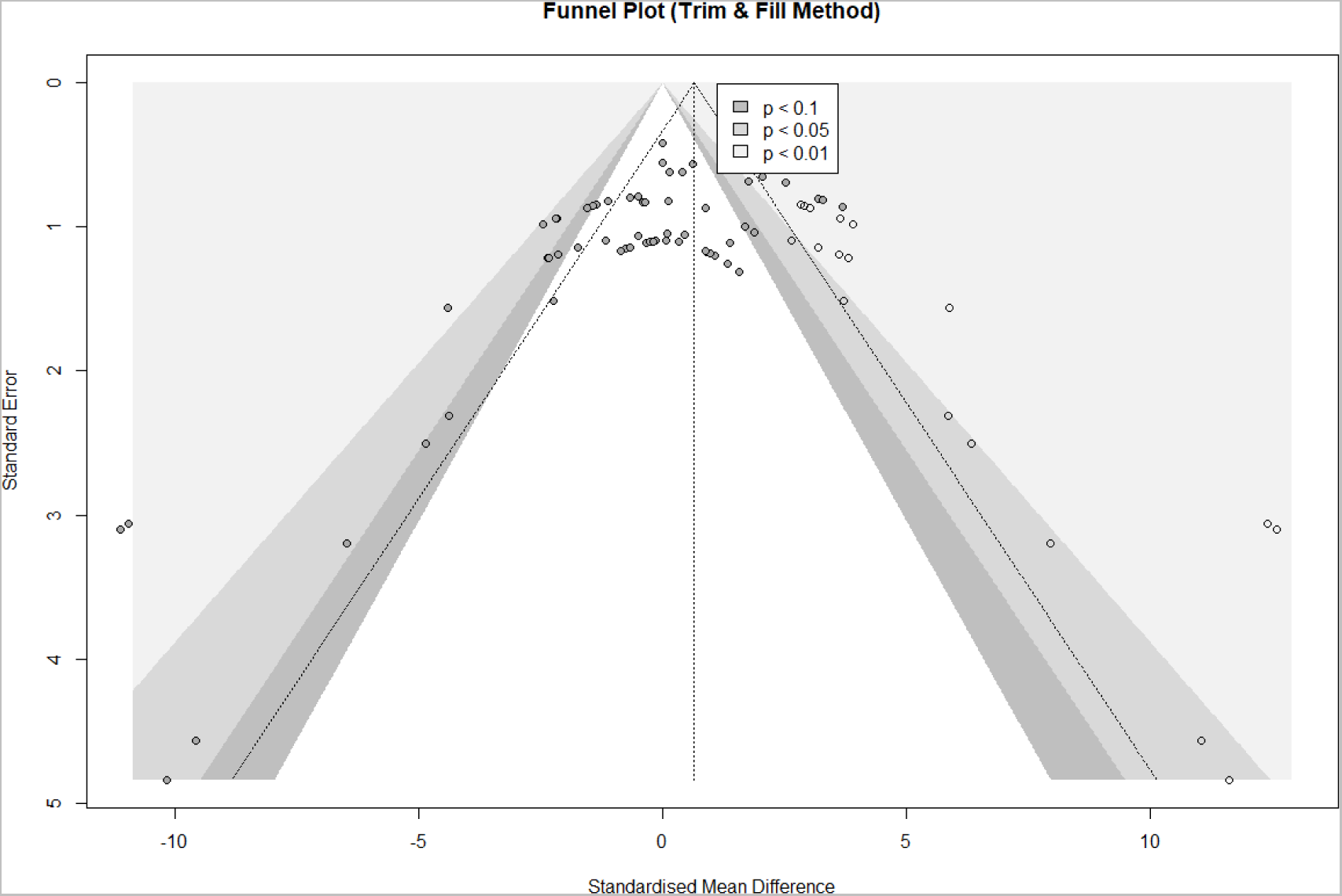
Funnel plot including studies analyzed within the outcome of spontaneous movements. Each gray circle represents a single comparison. Hollow circles represent imputed studies in the trim and fill analysis. The vertical line represents the overall effect size and the triangular region represents the 95% confidence interval. Shaded areas represent the interval for statistically significant effects.

Egger’s regression test indicated publication bias only for spontaneous movements, which showed a p < 0.0001 (for distance, p = 0.4120) (Table S1).

### 3.6) Sensitivity analysis

The leave-one-out analysis for distance revealed that none of the comparisons significantly modifies the result of the meta-analysis (Fig. 8). The overall effect and heterogeneity remained close to the original value. However, in order to confirm that any isolated study is skewing the results, we performed another meta-analysis excluding all the comparisons from the study by Li et al., 2019. In the forest plot, this study showed unusually high SMD, and the omission of their experiments in the leave-one-out analysis altered the overall effect direction. The significant overall effect was sustained (SMD −0.31 [−0.54; −0.08] (Fig. S8).

**Fig. 8.**
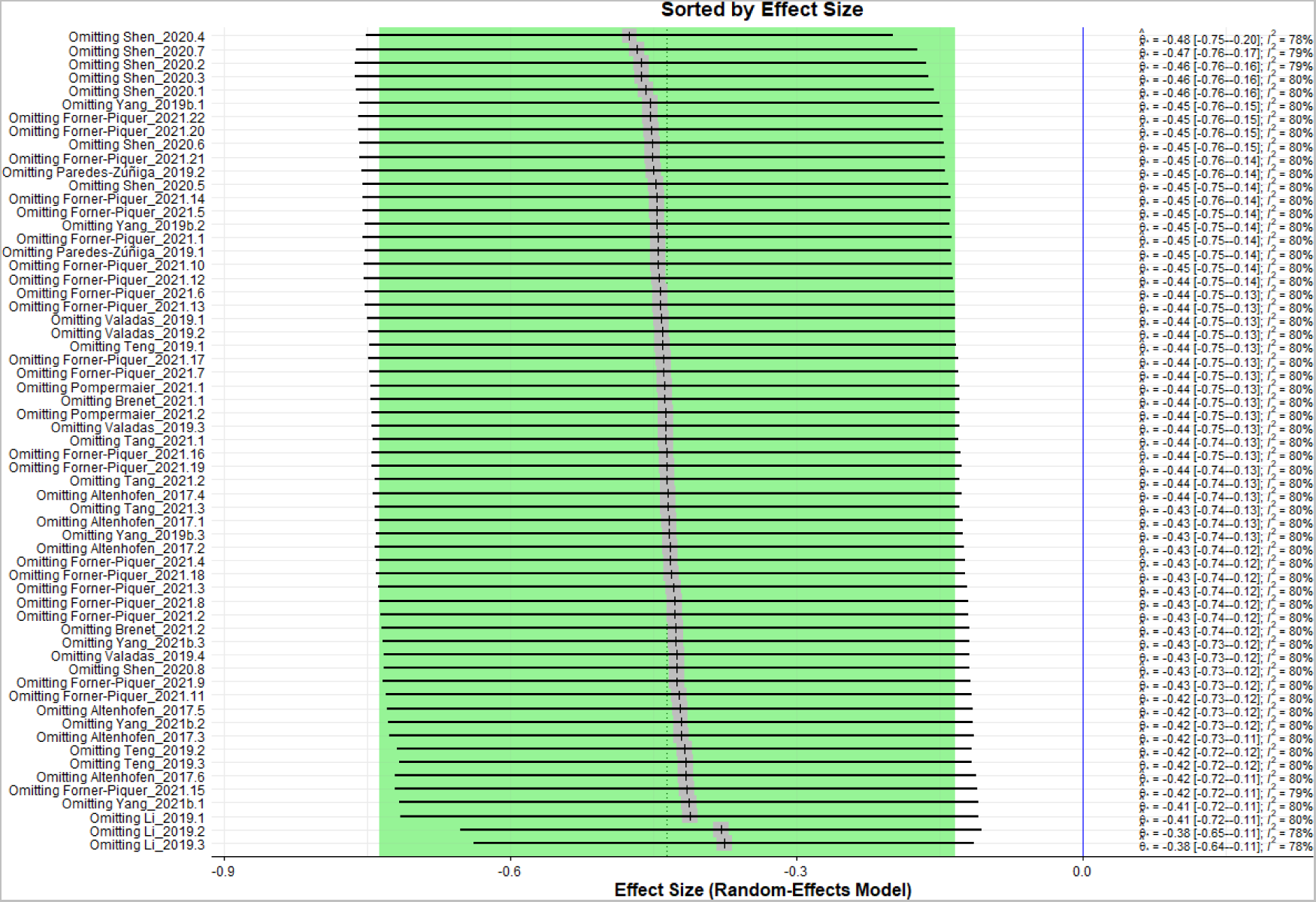
Sensitivity analyses for studies for the outcome of distance. Data are presented as Hedges’ G standardized mean differences (SMD) and 95% confidence intervals.

The leave-on-out analysis for spontaneous movements showed that omitting comparisons did not significantly modify the meta-analysis original result (Fig. 9). We also ran the meta-analysis without 2 studies: Cao et al., 2019b and Yang et al., 2016. In the forest plot, these studies showed atypically high SMD, and the omission of their experiments in the leave-one-out analysis changed the overall effect direction. Although the direction of the effect changed, it was still not significant (SMD 0.22 [−021; 066]) (Fig. S9).

**Fig. 9.**
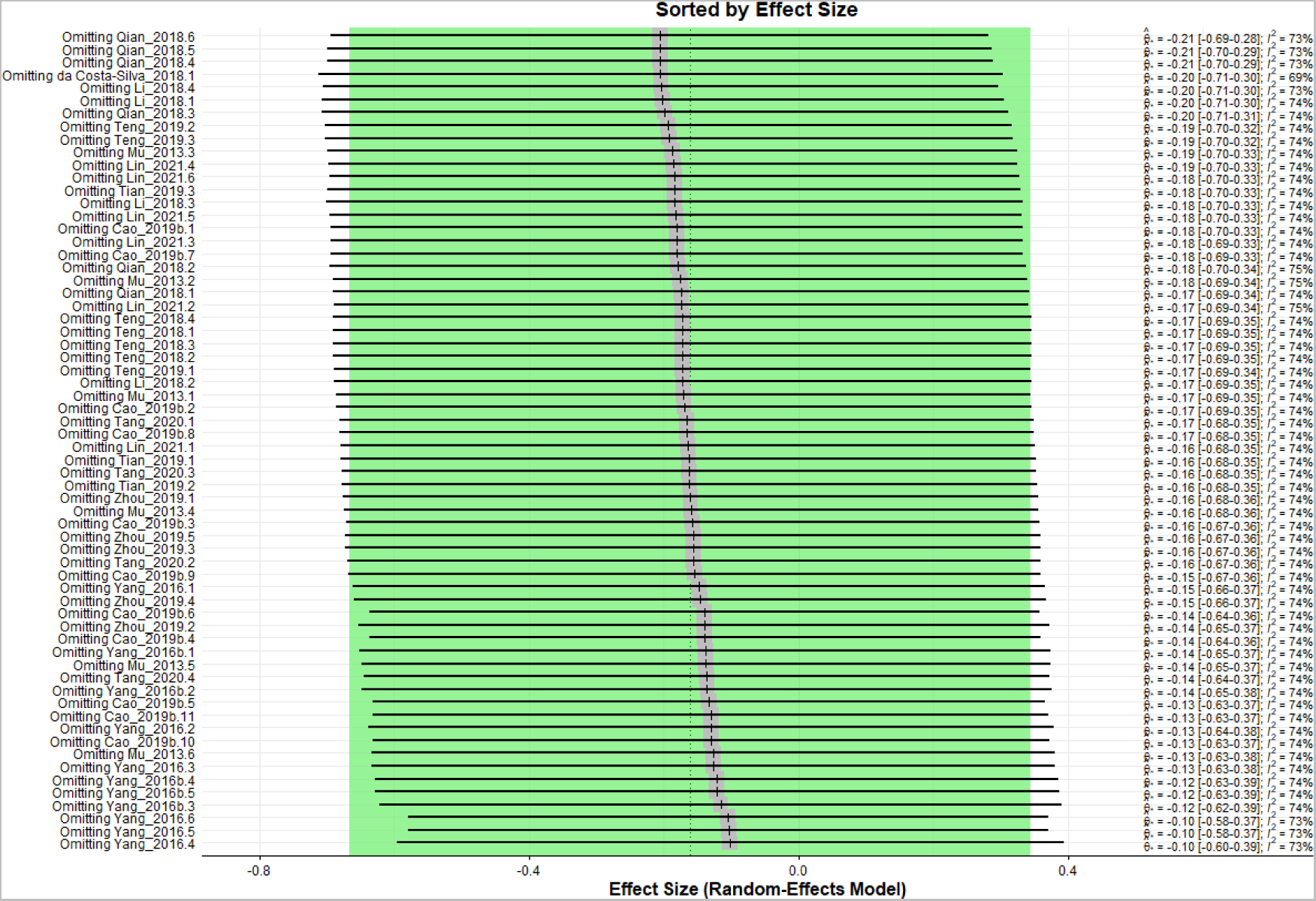
Sensitivity analyses for studies for the outcome of spontaneous movements. Data are presented as Hedges’ G standardized mean differences (SMD) and 95% confidence intervals.

## 4. Discussion

This work aimed to evaluate and synthesize the neurobehavioral effects of the exposure to fungicides in zebrafish through a systematic review and meta-analysis. We can highlight, as main findings, that fungicides cause a decrease in distance traveled by larval and adult zebrafish; no effect was observed on spontaneous movements of embryos.

Although there is no publication specifically synthesizing the effects of fungicide exposure in zebrafish, some articles approached this topic in a similar way. Review studies in the literature described the effects of pesticides as a whole, or isolated chemicals or classes of these compounds, using zebrafish or multiple species (Ames et al., 2022; Bhagat et al., 2021; Gonçalves et al., 2020; Lopes et al., 2022; Santana et al., 2021; Wang et al., 2021; Yanicostas and Soussi-Yanicostas, 2021; Zhang et al., 2020). Our results correlate with these previous findings in terms of neurotoxic effects and heterogeneity of the included studies.

The overall high heterogeneity observed in the meta-analysis for distance traveled can be attributed to several sources. The experimental conditions, from rearing until exposure and tests, are extremely variable between laboratories. The researchers employed a large number of protocols, which can include distinct durations of exposure, frequency of solution renewal, number of coexposed animals, age of the fish, type, and apparatus of the test (Hamm et al., 2019; Hill et al., 2023). When taking into account the subgroup analysis, included studies with adults had a low heterogeneity in comparison to those performed at the larval stage. Even though fewer adult studies were included, we can indeed verify more uniformity between the protocols of these experiments, mostly during the outcome assessment. Therefore, this similarity can explain the low heterogeneity of this subgroup.

Interestingly, there was no significant difference between the subgroups, indicating that the developmental stage of the animals does not significantly impact the effect of fungicides on distance traveled. Despite the different locomotor mechanisms exhibited by adults and larvae (Berg et al., 2018), it suggests that the exposure to fungicides affects both subgroups in a consistent manner.

On the other hand, the heterogeneity of the outcome of spontaneous movements was considered moderate. Unlike the distance traveled, the spontaneous movements can be measured in a single developmental stage: the embryo, generally at 24 hours post-fertilization (hpf). Consequently, the age of the animals can be excluded as a potential source of heterogeneity, which helps to explain why the heterogeneity did not reach the highest level.

The reporting quality analysis showed a high percentage of negative answers, especially relating to “sample size estimation” and “inclusion or exclusion criteria”. None of the authors explicitly stated previously determined parameters for the eligibility of the data. The result from this evaluation indicates that the conclusions of this review should be interpreted with caution, since the report of the included studies presents a considerable level of uncertainty. This lack of methodological information has been recognized as one of the main reasons behind the reproducibility crisis in preclinical research (Samsa and Samsa, 2019). Aiming to improve the quality of the studies, guidelines for the report of research with animals have been developed in the last years (Sert et al., 2020), however, it is a multifaceted problem that demands complex and long-term solutions (Begley and Ioannidis, 2015).

The results for the Egger’s test for publication bias were significant only for the outcome of spontaneous movements. There is an important role of selective publishing in the misinterpretation of a meta-analysis (Worp et al., 2010), which highlights the need for new practices regarding the publication of non-significant results.

The sensitivity analysis indicated that the results of the meta-analysis were not significantly influenced by any particular study or set of studies, suggesting that the overall effect size is robust and reliable. This finding supports the validity of the meta-analytic conclusions and can increase the confidence in the reliability of the results. However, the reliability of each individual comparison could not be determined due to poor reporting practices and a general lack of protocol preregistration.

## 5. Conclusion

Our results reinforce the toxicity of these chemicals, with their misuse representing a threat to the ecosystems. Since we depend on the affected environment, its contamination is an alert to public health. Besides that, we confirm the demand for well-designed studies with greater clarity of report on this topic, which allows more reliable conclusions aiming to contribute to a comprehensive monitoring of environmental pollution.

## 6. Data availability

All data are available at Open Science Framework (https://osf.io/hdu5c/).

## Authors contributions

**Carlos G. Reis**: conceptualization, data curation, formal analysis, investigation, methodology, project administration, visualization, and writing - original draft; **Leonardo M. Bastos**: conceptualization, investigation, methodology, visualization, and writing – review & editing; **Rafael Chitolina**: conceptualization, investigation, methodology, and writing – review & editing; **Matheus Gallas-Lopes**: data curation, formal analysis, methodology, visualization, and writing – review & editing; **Querusche K. Zanona**: conceptualization, methodology, and writing – review & editing; **Sofia Z. Becker**: conceptualization, investigation, and writing – review & editing; **Ana P. Herrmann**: conceptualization, investigation, methodology, supervision, project administration, and writing - review & editing; **Angelo Piato**: conceptualization, investigation, methodology, supervision, project administration, and writing – review & editing.

## Declaration of competing interest

The authors declare no competing interest.

## Acknowledgments

We thank the Conselho Nacional de Desenvolvimento Científico e Tecnológico (CNPq, proc. 303343/2020-6), Coordenação de Aperfeiçoamento de Pessoal de Nível Superior - Brasil (CAPES), and Pró-Reitoria de Pesquisa (PROPESQ) at Universidade Federal do Rio Grande do Sul (UFRGS) for funding and support.

## Funding

CGR is recipient of a fellowship from CAPES.

## Supplementary material

**Fig. S1.**
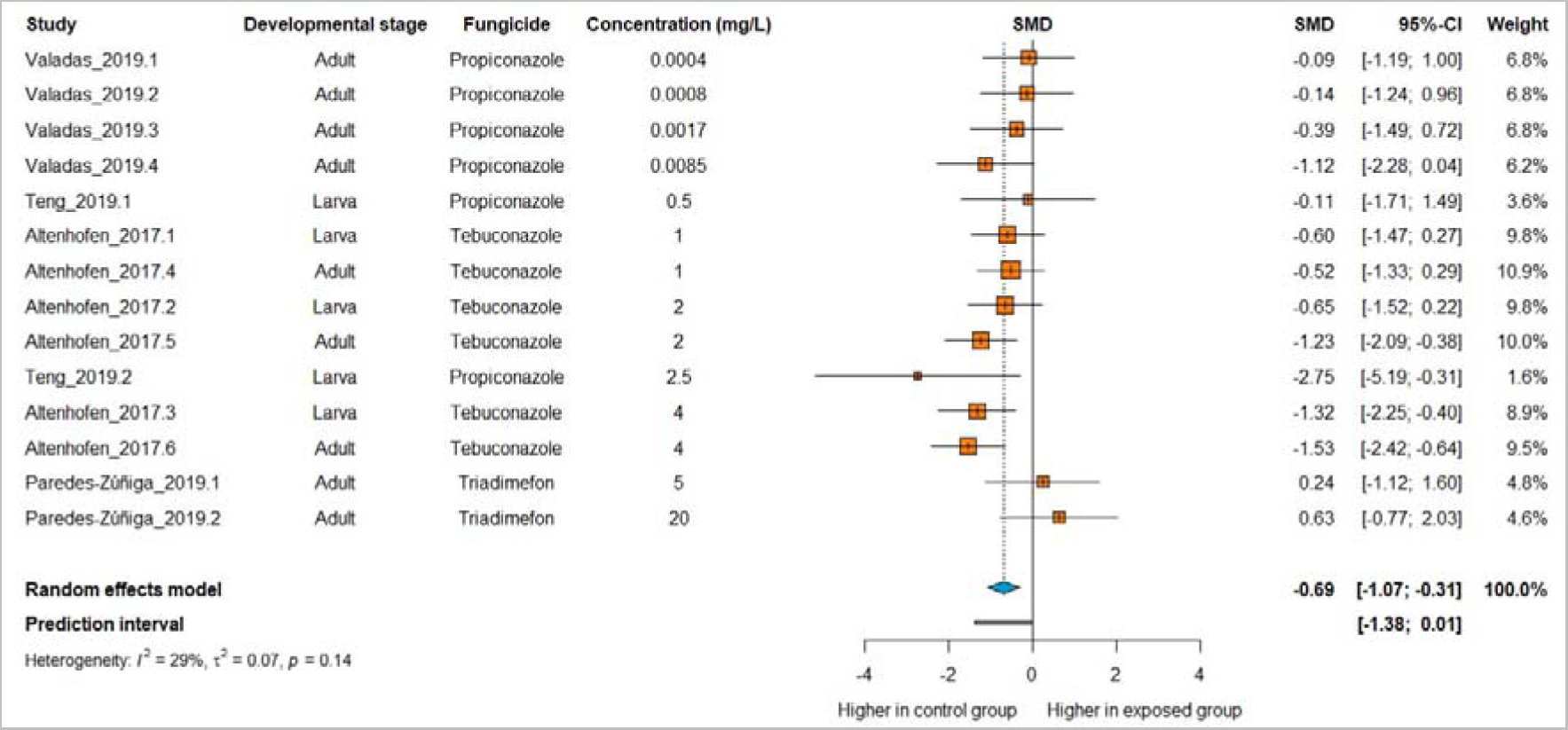
The effect of exposure to triazole fungicides on distance traveled in zebrafish. Data are presented as Hedges’ G standardized mean differences (SMD) and 95% confidence intervals.

**Fig. S2.**
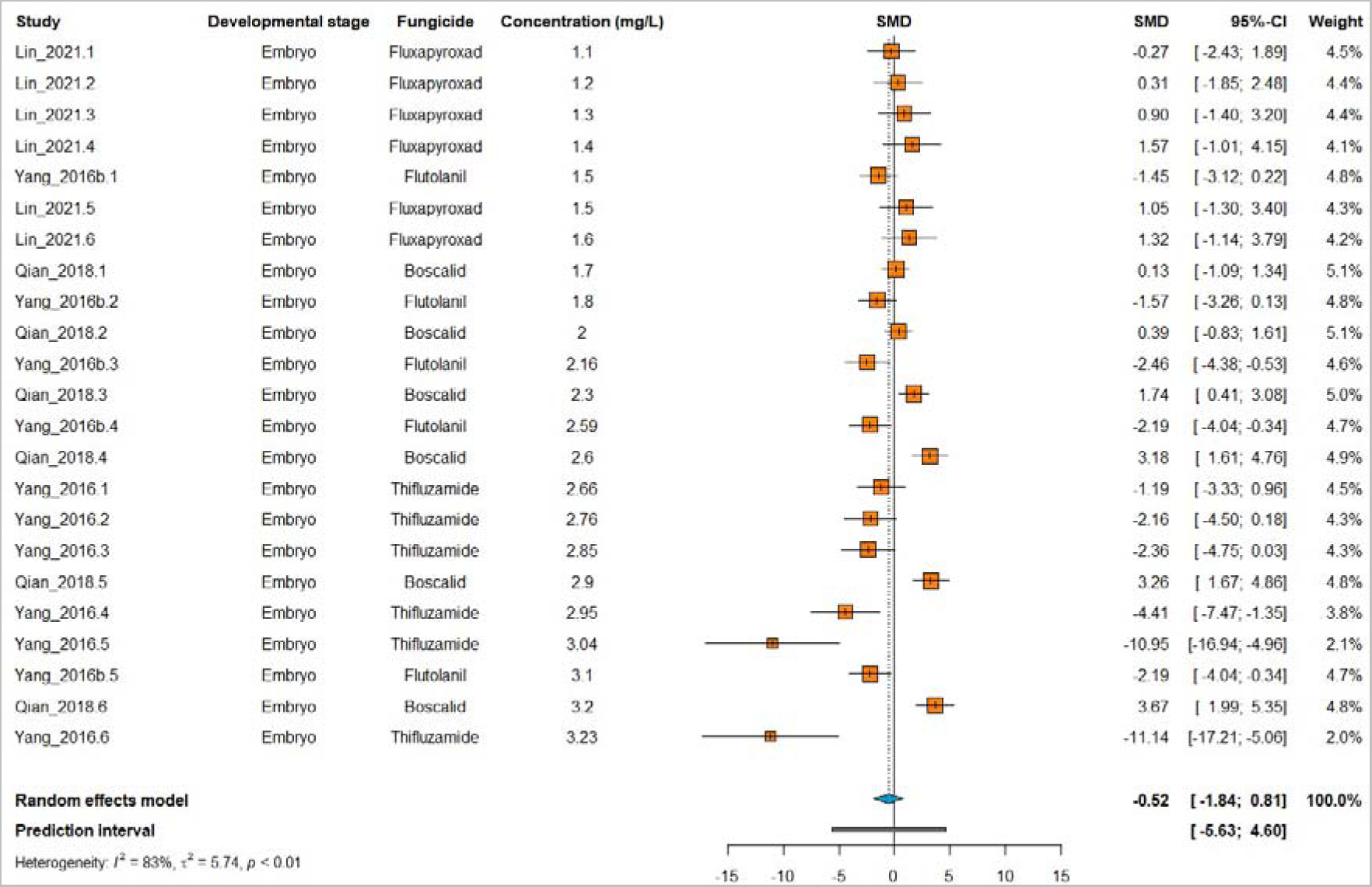
The effect of exposure to anilide fungicides on spontaneous movements in zebrafish. Data are presented as Hedges’ G standardized mean differences (SMD) and 95% confidence intervals.

**Fig. S3.**
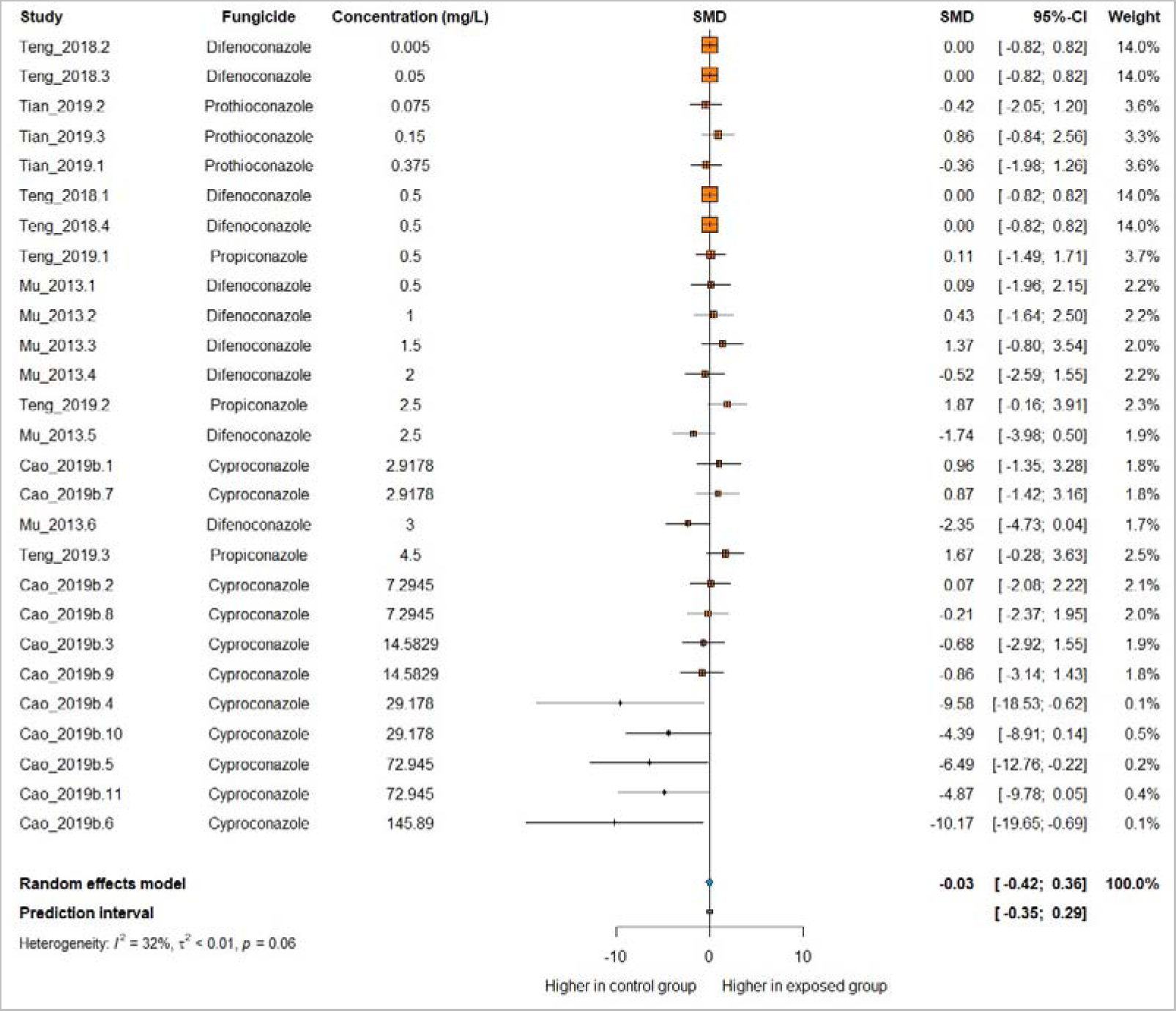
The effect of exposure to triazole fungicides on spontaneous movements in zebrafish. Data are presented as Hedges’ G standardized mean differences (SMD) and 95% confidence intervals.

**Fig. S4.**
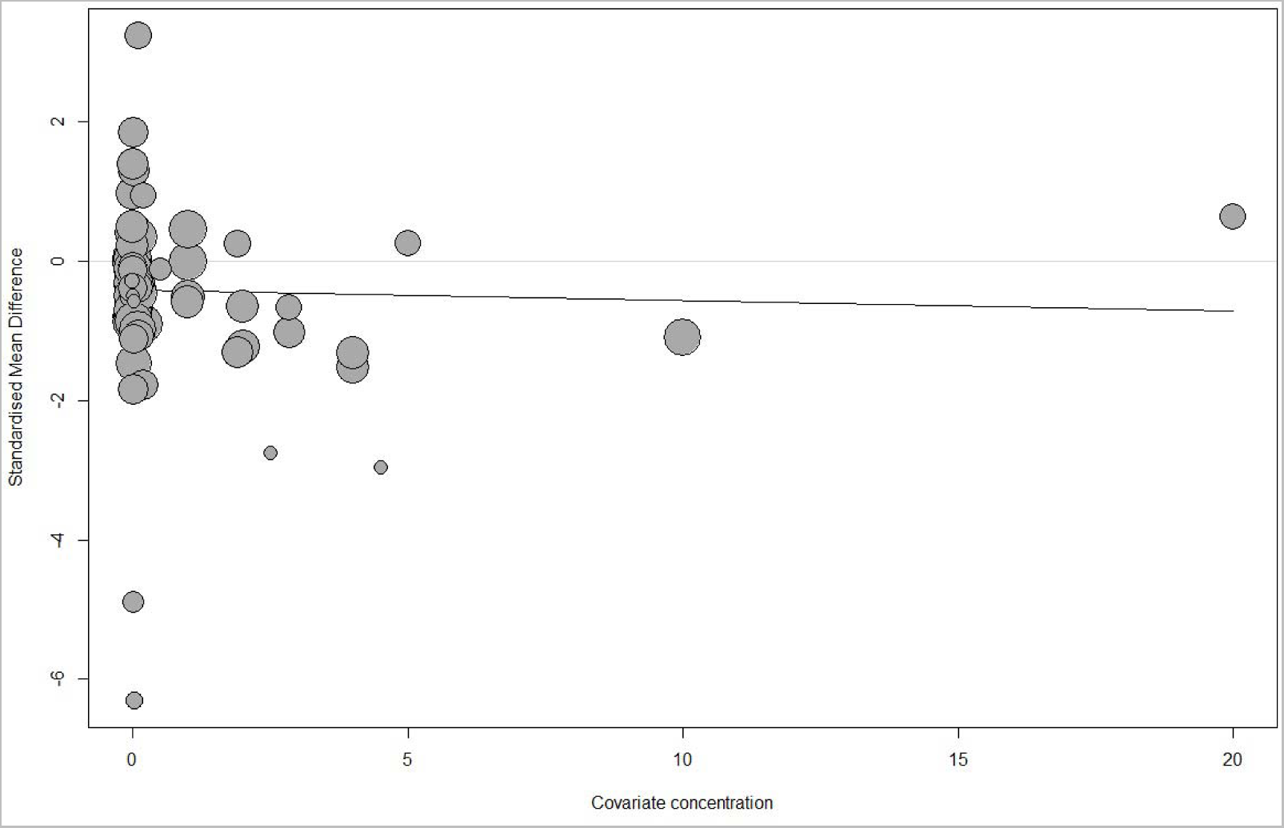
Meta-regression of distance using the concentration of fungicides as the moderator variable.

**Fig. S5.**
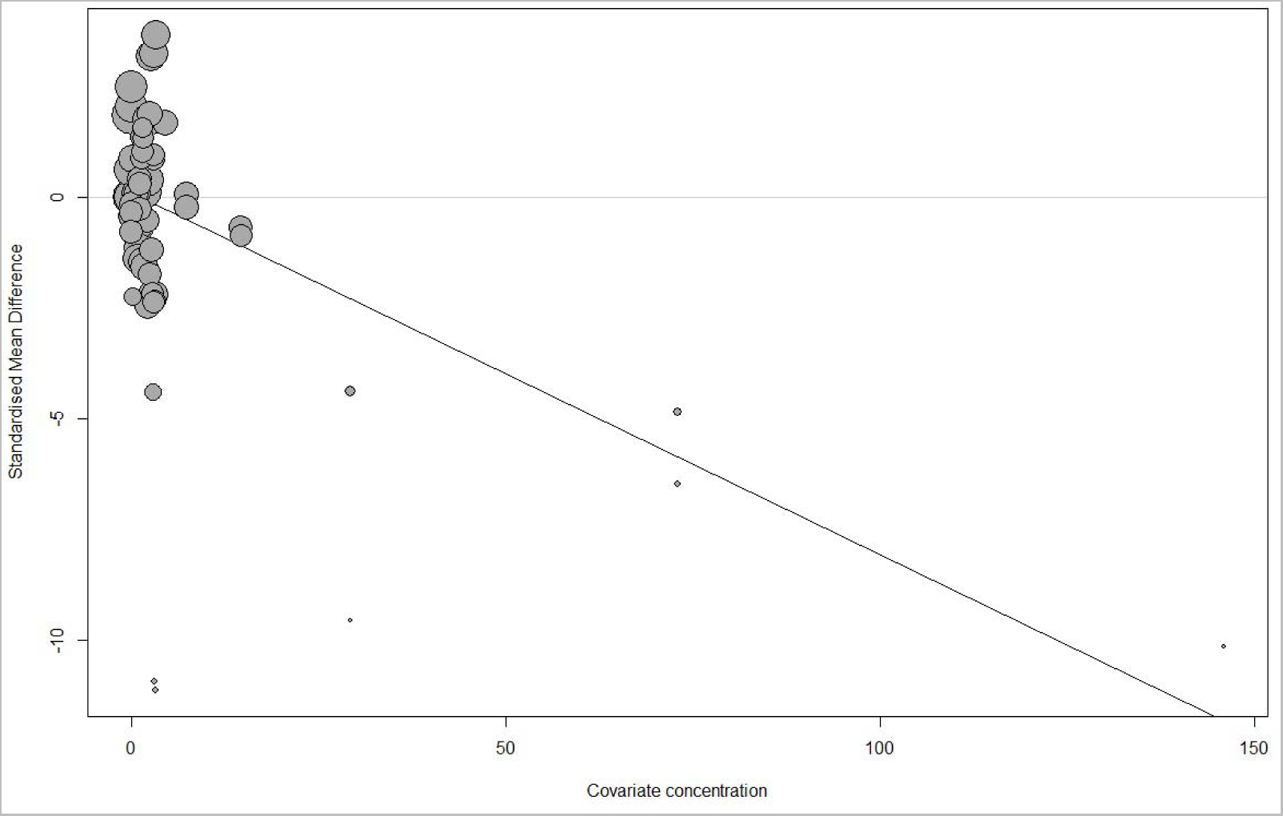
Meta-regression of spontaneous movements using the concentration of fungicides as the moderator variable.

**Fig. S6.**
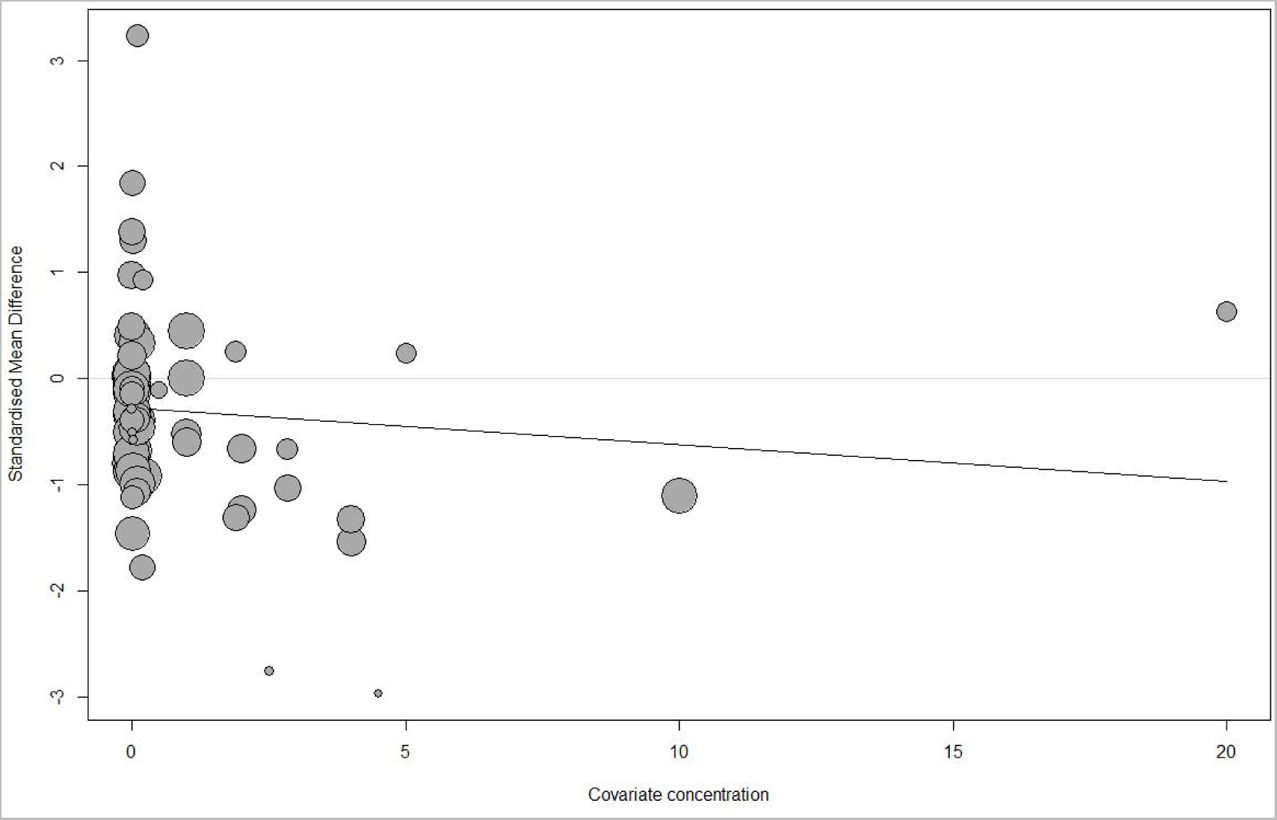
Meta-regression of distance excluding the study by Li, 2019 using the concentration of fungicide as the moderator variable.

**Fig. S7.**
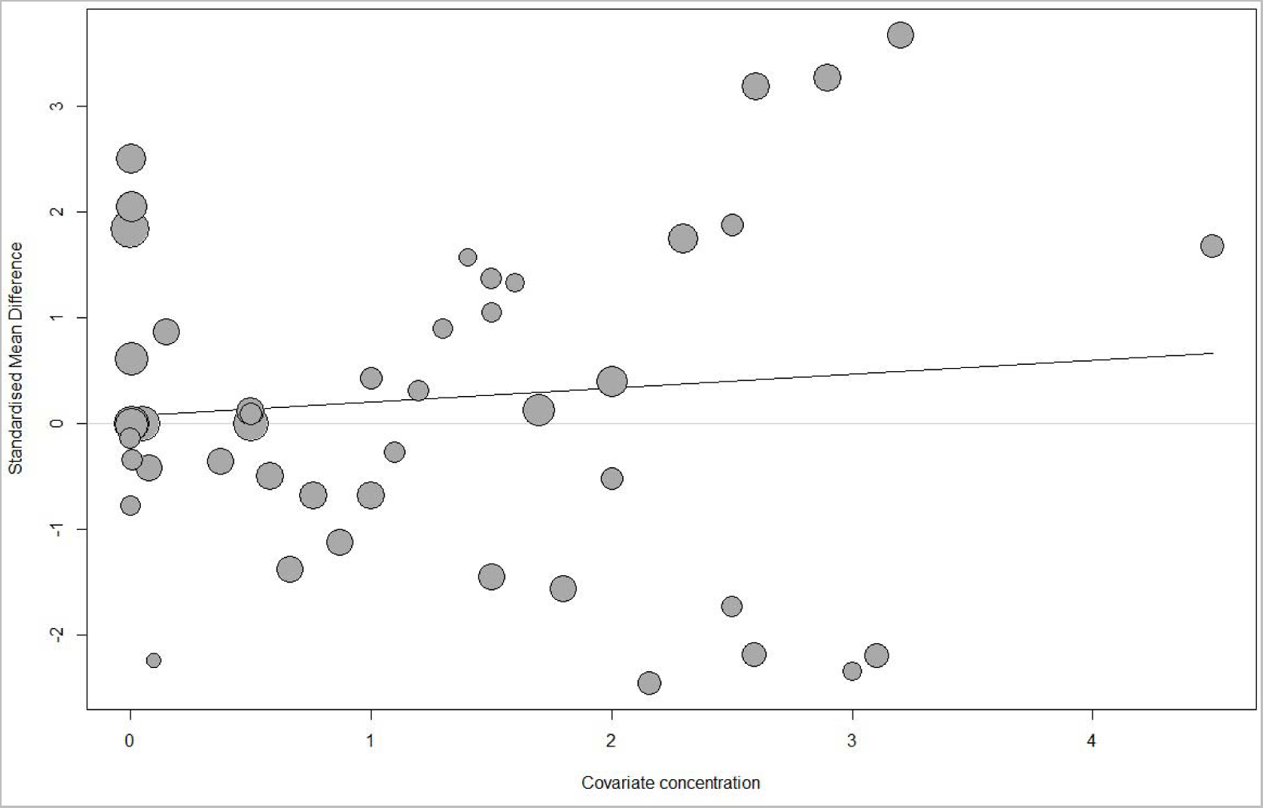
Meta-regression of spontaneous movements excluding the studies by Cao, 2019b and Yang, 2016 using the concentration of fungicide as the moderator variable.

**Table S1.** Egger’s regression test summary for the outcomes of distance and spontaneous movements.

| **Outcome** | **Intercept** | **Standard error** | **t** | **p-value** |
| --- | --- | --- | --- | --- |
| Distance | -0.2221 | 0.1990 | -0.83 | 0.4120 |
| Spontaneous movements | 1.8723 | 0.3385 | -5.10 | < 0.0001 |

**Fig. S8.**
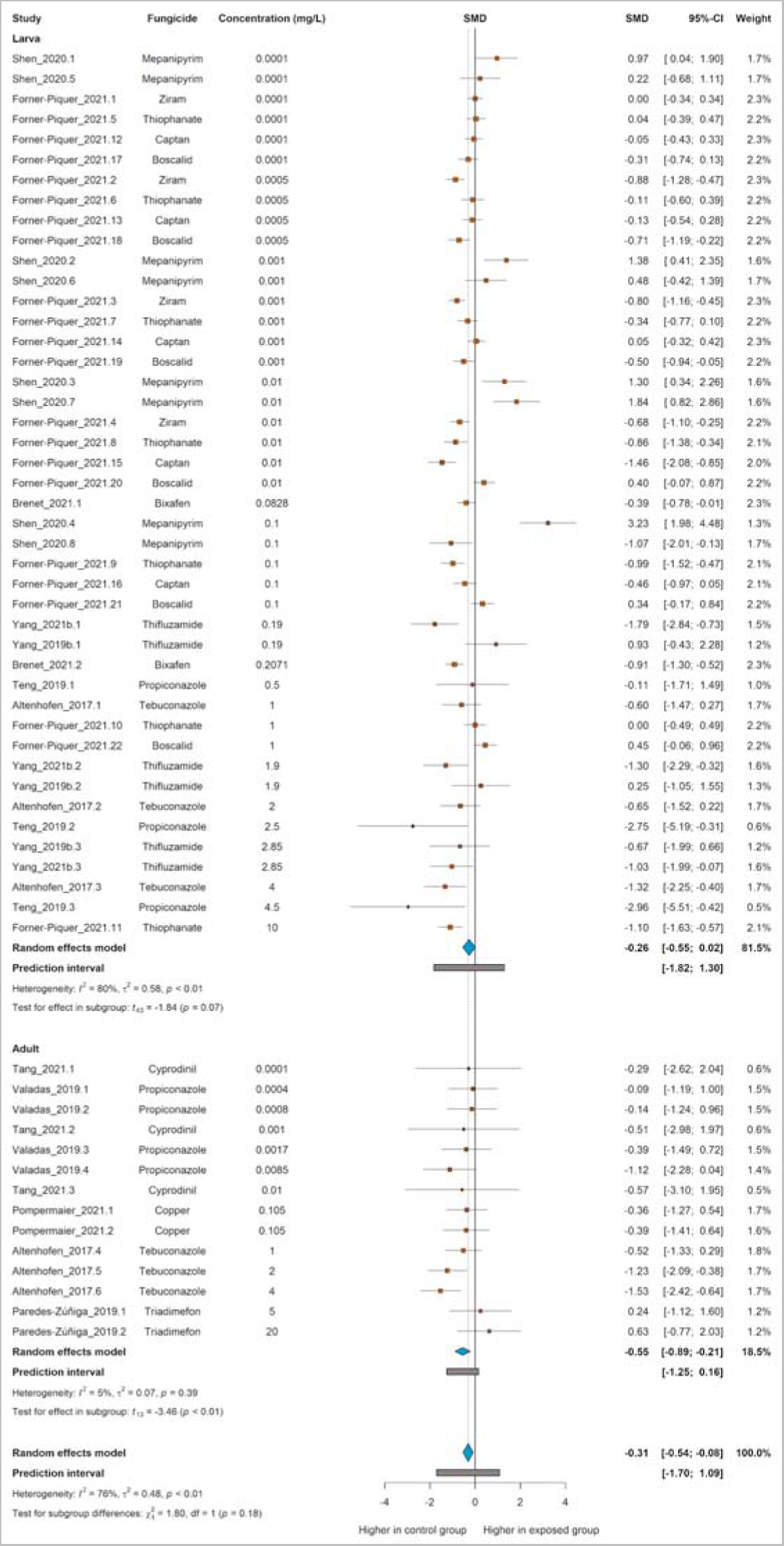
The effect of exposure to fungicides on distance traveled in zebrafish excluding the study by Li, 2019. Data are presented as Hedges’ G standardized mean differences (SMD) and 95% confidence intervals.

**Fig. S9.**
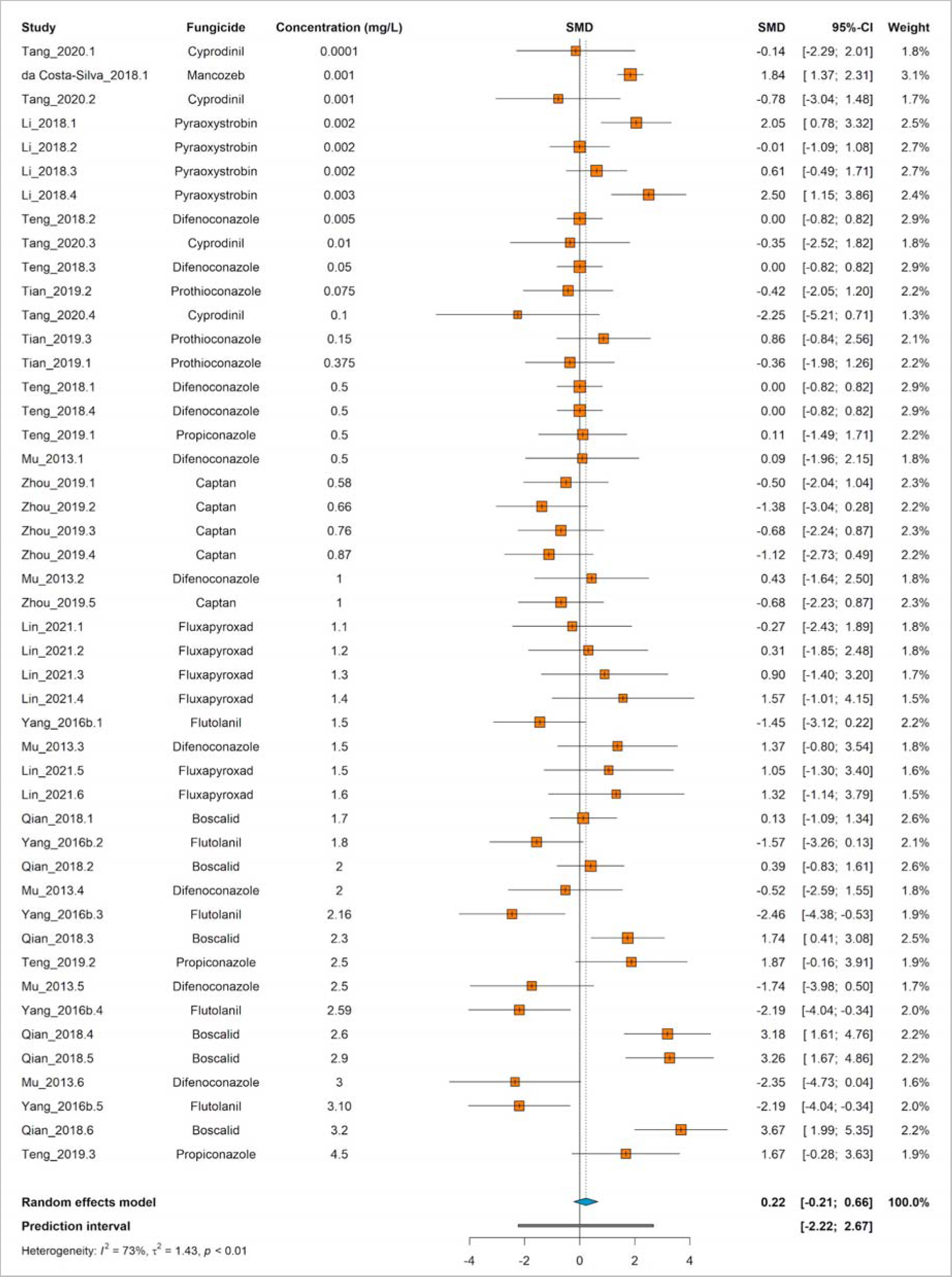
The effect of exposure to fungicides on spontaneous movements in zebrafish excluding the studies by Cao, 2019b and Yang, 2016. Data are presented as Hedges’ G standardized mean differences (SMD) and 95% confidence intervals.

## Notes

### Competing Interest Statement

The authors have declared no competing interest.

## References

AERU, 2022. Pesticide Properties Database. Agriculture and Environment Research Unit (AERU) at the University of Hertfordshire. [WWW Document]. URL http://sitem.herts.ac.uk/aeru/ppdb/en/atoz_fung.htm (accessed 5.15.23).

Akoto, O., Azuure, A.A., Adotey, K.D., 2016. Pesticide residues in water, sediment and fish from Tono Reservoir and their health risk implications. SpringerPlus 5, 1849. https://doi.org/10.1186/s40064-016-3544-z

Albuquerque, A.F., Ribeiro, J.S., Kummrow, F., Nogueira, A.J.A., Montagner, C.C., Umbuzeiro, G.A., 2016. Pesticides in Brazilian freshwaters: a critical review. Environ. Sci. Process. Impacts 18, 779–787. https://doi.org/10.1039/c6em00268d

Altenhofen, S., Nabinger, D.D., Wiprich, M.T., Pereira, T.C.B., Bogo, M.R., Bonan, C.D., 2017. Tebuconazole alters morphological, behavioral and neurochemical parameters in larvae and adult zebrafish (Danio rerio). Chemosphere 180, 483–490. https://doi.org/10.1016/j.chemosphere.2017.04.029

Ames, J., Miragem, A.A., Cordeiro, M.F., Cerezer, F.O., Loro, V.L., 2022. Effects of glyphosate on zebrafish: a systematic review and meta-analysis. Ecotoxicol. Lond. Engl. 31, 1189–1204. https://doi.org/10.1007/s10646-022-02581-z

Andrade, C., 2020. Mean Difference, Standardized Mean Difference (SMD), and Their Use in Meta-Analysis: As Simple as It Gets. J. Clin. Psychiatry 81, 20f13681. https://doi.org/10.4088/JCP.20f13681

Andrade, T.S., Henriques, J.F., Almeida, A.R., Machado, A.L., Koba, O., Giang, P.T., Soares, A.M.V.M., Domingues, I., 2016. Carbendazim exposure induces developmental, biochemical and behavioural disturbance in zebrafish embryos. Aquat. Toxicol. Amst. Neth. 170, 390–399. https://doi.org/10.1016/j.aquatox.2015.11.017

Bakkers, J., 2011. Zebrafish as a model to study cardiac development and human cardiac disease. Cardiovasc. Res. 91, 279–288. https://doi.org/10.1093/cvr/cvr098

Balduzzi, S., Rücker, G., Schwarzer, G., 2019. How to perform a meta-analysis with R: a practical tutorial. Evid. Based Ment. Health 22, 153–160. https://doi.org/10.1136/ebmental-2019-300117

Barreto, A., Santos, J., Amorim, M.J.B., Maria, V.L., 2021. Is the Synthetic Fungicide Fosetyl-Al Safe for the Ecotoxicological Models Danio rerio and Enchytraeus crypticus? Appl. Sci. 11, 7209. https://doi.org/10.3390/app11167209

Beckerman, J., Palmer, C., Tedford, E., Ypema, H., 2023. Fifty Years of Fungicide Development, Deployment, and Future Use. Phytopathology® 113, 694–706. https://doi.org/10.1094/PHYTO-10-22-0399-IA

Begley, C.G., Ioannidis, J.P.A., 2015. Reproducibility in science: improving the standard for basic and preclinical research. Circ. Res. 116, 116–126. https://doi.org/10.1161/CIRCRESAHA.114.303819

Beketov, M.A., Kefford, B.J., Schäfer, R.B., Liess, M., 2013. Pesticides reduce regional biodiversity of stream invertebrates. Proc. Natl. Acad. Sci. U. S. A. 110, 11039–11043. https://doi.org/10.1073/pnas.1305618110

Berg, E.M., Björnfors, E.R., Pallucchi, I., Picton, L.D., El Manira, A., 2018. Principles Governing Locomotion in Vertebrates: Lessons From Zebrafish. Front. Neural Circuits 12, 73. https://doi.org/10.3389/fncir.2018.00073

Bhagat, J., Singh, N., Nishimura, N., Shimada, Y., 2021. A comprehensive review on environmental toxicity of azole compounds to fish. Chemosphere 262, 128335. https://doi.org/10.1016/j.chemosphere.2020.128335

Brenet, A., Hassan-Abdi, R., Soussi-Yanicostas, N., 2021. Bixafen, a succinate dehydrogenase inhibitor fungicide, causes microcephaly and motor neuron axon defects during development. Chemosphere 265, 128781. https://doi.org/10.1016/j.chemosphere.2020.128781

Cao, F., Souders, C.L., Li, P., Adamovsky, O., Pang, S., Qiu, L., Martyniuk, C.J., 2019a. Developmental toxicity of the fungicide ziram in zebrafish (Danio rerio). Chemosphere 214, 303–313. https://doi.org/10.1016/j.chemosphere.2018.09.105

Cao, F., Souders, C.L., Li, P., Pang, S., Qiu, L., Martyniuk, C.J., 2019b. Developmental toxicity of the triazole fungicide cyproconazole in embryo-larval stages of zebrafish (Danio rerio). Environ. Sci. Pollut. Res. 26, 4913–4923. https://doi.org/10.1007/s11356-018-3957-z

Cao, F., Souders, C.L., Li, P., Pang, S., Liang, X., Qiu, L., Martyniuk, C.J., 2019c. Developmental neurotoxicity of maneb: Notochord defects, mitochondrial dysfunction and hypoactivity in zebrafish (Danio rerio) embryos and larvae. Ecotoxicol. Environ. Saf. 170, 227–237. https://doi.org/10.1016/j.ecoenv.2018.11.110

Clasen, B., Loro, V.L., Murussi, C.R., Tiecher, T.L., Moraes, B., Zanella, R., 2018. Bioaccumulation and oxidative stress caused by pesticides in Cyprinus carpio reared in a rice-fish system. Sci. Total Environ. 626, 737–743. https://doi.org/10.1016/j.scitotenv.2018.01.154

Cochran, W.G., 1954. Some Methods for Strengthening the Common χ2 Tests. Biometrics 10, 417–451. https://doi.org/10.2307/3001616

Costa-Silva, D.G. da, Leandro, L.P., Vieira, P. de B., de Carvalho, N.R., Lopes, A.R., Schimith, L.E., Nunes, M.E.M., de Mello, R.S., Martins, I.K., de Paula, A.A., Cañedo, A.D., Moreira, J.C.F., Posser, T., Franco, J.L., 2018. N-acetylcysteine inhibits Mancozeb-induced impairments to the normal development of zebrafish embryos. Neurotoxicol. Teratol. 68, 1–12. https://doi.org/10.1016/j.ntt.2018.04.003

Dai, Y.-J., Jia, Y.-F., Chen, N., Bian, W.-P., Li, Q.-K., Ma, Y.-B., Chen, Y.-L., Pei, D.-S., 2014. Zebrafish as a model system to study toxicology. Environ. Toxicol. Chem. 33, 11–17. https://doi.org/10.1002/etc.2406

de Araújo, E.P., Caldas, E.D., Oliveira-Filho, E.C., 2022. Pesticides in surface freshwater: a critical review. Environ. Monit. Assess. 194, 452. https://doi.org/10.1007/s10661-022-10005-y

De la Paz, J.F., Beiza, N., Paredes-Zúñiga, S., Hoare, M.S., Allende, M.L., 2017. Triazole Fungicides Inhibit Zebrafish Hatching by Blocking the Secretory Function of Hatching Gland Cells. Int. J. Mol. Sci. 18, 710. https://doi.org/10.3390/ijms18040710

Delgado-Blanca, I., Ruiz-Medina, A., Ortega-Barrales, P., 2019. Novel sequential separation and determination of a quaternary mixture of fungicides by using an automatic fluorimetric optosensor. Food Addit. Contam. Part Chem. Anal. Control Expo. Risk Assess. 36, 278–288. https://doi.org/10.1080/19440049.2018.1564372

Domingues, I., Oliveira, R., Musso, C., Cardoso, M., Soares, A.M.V.M., Loureiro, S., 2013. Prochloraz effects on biomarkers activity in zebrafish early life stages and adults. Environ. Toxicol. 28, 155–163. https://doi.org/10.1002/tox.20710

Duval, S., Tweedie, R., 2000. Trim and fill: A simple funnel-plot-based method of testing and adjusting for publication bias in meta-analysis. Biometrics 56, 455–463. https://doi.org/10.1111/j.0006-341x.2000.00455.x

Egger, M., Davey Smith, G., Schneider, M., Minder, C., 1997. Bias in meta-analysis detected by a simple, graphical test. BMJ 315, 629–634. https://doi.org/10.1136/bmj.315.7109.629

Fan, R., Zhang, W., Li, L., Jia, L., Zhao, J., Zhao, Z., Peng, S., Yuan, X., Chen, Y., 2021. Individual and synergistic toxic effects of carbendazim and chlorpyrifos on zebrafish embryonic development. Chemosphere 280, 130769. https://doi.org/10.1016/j.chemosphere.2021.130769

Fan, Y., Miao, W., Lai, K., Huang, W., Song, R., Li, Q.X., 2018. Developmental toxicity and inhibition of the fungicide hymexazol to melanin biosynthesis in zebrafish embryos. Pestic. Biochem. Physiol., Special Issue: Fungicide Toxicology in China 147, 139–144. https://doi.org/10.1016/j.pestbp.2017.10.007

FAO, WHO, 2022. Manual on the development and use of FAO and WHO specifications for chemical pesticides. FAO, WHO, Rome, Italy. https://doi.org/10.4060/cb8401en

FAOSTAT, 2023. Statistics Division of the Food and Agriculture Organization of the United Nations [WWW Document]. URL https://www.fao.org/faostat/en/#data/RP/visualize (accessed 5.17.23).

Fitzgerald, J.A., Könemann, S., Krümpelmann, L., Županič, A., Vom Berg, C., 2021. Approaches to Test the Neurotoxicity of Environmental Contaminants in the Zebrafish Model: From Behavior to Molecular Mechanisms. Environ. Toxicol. Chem. 40, 989–1006. https://doi.org/10.1002/etc.4951

Fitzmaurice, A.G., Rhodes, S.L., Lulla, A., Murphy, N.P., Lam, H.A., O’Donnell, K.C., Barnhill, L., Casida, J.E., Cockburn, M., Sagasti, A., Stahl, M.C., Maidment, N.T., Ritz, B., Bronstein, J.M., 2013. Aldehyde dehydrogenase inhibition as a pathogenic mechanism in Parkinson disease. Proc. Natl. Acad. Sci. 110, 636– 641. https://doi.org/10.1073/pnas.1220399110

Forner-Piquer, I., Klement, W., Gangarossa, G., Zub, E., de Bock, F., Blaquiere, M., Maurice, T., Audinat, E., Faucherre, A., Lasserre, F., Ellero-Simatos, S., Gamet-Payrastre, L., Jopling, C., Marchi, N., 2021. Varying modalities of perinatal exposure to a pesticide cocktail elicit neurological adaptations in mice and zebrafish. Environ. Pollut. 278, 116755. https://doi.org/10.1016/j.envpol.2021.116755

Frommlet, F., 2020. Improving reproducibility in animal research. Sci. Rep. 10, 19239. https://doi.org/10.1038/s41598-020-76398-3

Gerlai, R., 2019. Reproducibility and replicability in zebrafish behavioral neuroscience research. Pharmacol. Biochem. Behav. 178, 30–38. https://doi.org/10.1016/j.pbb.2018.02.005

Gonçalves, Í.F.S., Souza, T.M., Vieira, L.R., Marchi, F.C., Nascimento, A.P., Farias, D.F., 2020. Toxicity testing of pesticides in zebrafish—a systematic review on chemicals and associated toxicological endpoints. Environ. Sci. Pollut. Res. 27, 10185–10204. https://doi.org/10.1007/s11356-020-07902-5

Hamm, J.T., Ceger, P., Allen, D., Stout, M., Maull, E.A., Baker, G., Zmarowski, A., Padilla, S., Perkins, E., Planchart, A., Stedman, D., Tal, T., Tanguay, R.L., Volz, D.C., Wilbanks, M.S., Walker, N.J., 2019. Characterizing sources of variability in zebrafish embryo screening protocols. ALTEX 36, 103–120. https://doi.org/10.14573/altex.1804162

Higgins, J.P.T., Thomas, J., Chandler, J., Cumpston, M., Li, T., Page, M.J., Welch, V.A., 2019. Cochrane handbook for systematic reviews of interventions. John Wiley & Sons.

Higgins, J.P.T., Thompson, S.G., 2002. Quantifying heterogeneity in a meta-analysis. Stat. Med. 21, 1539–1558. https://doi.org/10.1002/sim.1186

Hill, B.N., Britton, K.N., Hunter, D.L., Olin, J.K., Lowery, M., Hedge, J.M., Knapp, B.R., Jarema, K.A., Rowson, Z., Padilla, S., 2023. Inconsistencies in variable reporting and methods in larval zebrafish behavioral assays. Neurotoxicol. Teratol. 96, 107163. https://doi.org/10.1016/j.ntt.2023.107163

Howe, K., Clark, M.D., Torroja, C.F., Torrance, J., Berthelot, C., Muffato, M., Collins, J.E., Humphray, S., McLaren, K., Matthews, L., McLaren, S., Sealy, I., Caccamo, M., Churcher, C., Scott, C., Barrett, J.C., Koch, R., Rauch, G.-J., White, S., Chow, W., Kilian, B., Quintais, L.T., Guerra-Assunção, J.A., Zhou, Y., Gu, Y., Yen, J., Vogel, J.-H., Eyre, T., Redmond, S., Banerjee, R., Chi, J., Fu, B., Langley, E., Maguire, S.F., Laird, G.K., Lloyd, D., Kenyon, E., Donaldson, S., Sehra, H., Almeida-King, J., Loveland, J., Trevanion, S., Jones, M., Quail, M., Willey, D., Hunt, A., Burton, J., Sims, S., McLay, K., Plumb, B., Davis, J., Clee, C., Oliver, K., Clark, R., Riddle, C., Elliot, D., Eliott, D., Threadgold, G., Harden, G., Ware, D., Begum, S., Mortimore, B., Mortimer, B., Kerry, G., Heath, P., Phillimore, B., Tracey, A., Corby, N., Dunn, M., Johnson, C., Wood, J., Clark, S., Pelan, S., Griffiths, G., Smith, M., Glithero, R., Howden, P., Barker, N., Lloyd, C., Stevens, C., Harley, J., Holt, K., Panagiotidis, G., Lovell, J., Beasley, H., Henderson, C., Gordon, D., Auger, K., Wright, D., Collins, J., Raisen, C., Dyer, L., Leung, K., Robertson, L., Ambridge, K., Leongamornlert, D., McGuire, S., Gilderthorp, R., Griffiths, C., Manthravadi, D., Nichol, S., Barker, G., Whitehead, S., Kay, M., Brown, J., Murnane, C., Gray, E., Humphries, M., Sycamore, N., Barker, D., Saunders, D., Wallis, J., Babbage, A., Hammond, S., Mashreghi-Mohammadi, M., Barr, L., Martin, S., Wray, P., Ellington, A., Matthews, N., Ellwood, M., Woodmansey, R., Clark, G., Cooper, J.D., Cooper, J., Tromans, A., Grafham, D., Skuce, C., Pandian, R., Andrews, R., Harrison, E., Kimberley, A., Garnett, J., Fosker, N., Hall, R., Garner, P., Kelly, D., Bird, C., Palmer, S., Gehring, I., Berger, A., Dooley, C.M., Ersan-Ürün, Z., Eser, C., Geiger, H., Geisler, M., Karotki, L., Kirn, A., Konantz, J., Konantz, M., Oberländer, M., Rudolph-Geiger, S., Teucke, M., Lanz, C., Raddatz, G., Osoegawa, K., Zhu, B., Rapp, A., Widaa, S., Langford, C., Yang, F., Schuster, S.C., Carter, N.P., Harrow, J., Ning, Z., Herrero, J., Searle, S.M.J., Enright, A., Geisler, R., Plasterk, R.H.A., Lee, C., Westerfield, M., de Jong, P.J., Zon, L.I., Postlethwait, J.H., Nüsslein-Volhard, C., Hubbard, T.J.P., Roest Crollius, H., Rogers, J., Stemple, D.L., 2013. The zebrafish reference genome sequence and its relationship to the human genome. Nature 496, 498–503. https://doi.org/10.1038/nature12111

Huang, T., Souders, C.L., Wang, S., Ganter, J., He, J., Zhao, Y.H., Cheng, H., Martyniuk, C.J., 2021. Behavioral and developmental toxicity assessment of the strobilurin fungicide fenamidone in zebrafish embryos/larvae (Danio rerio). Ecotoxicol. Environ. Saf. 228, 112966. https://doi.org/10.1016/j.ecoenv.2021.112966

Hussain, A., Audira, G., Malhotra, N., Uapipatanakul, B., Chen, J.-R., Lai, Y.- H., Huang, J.-C., Chen, K.H.-C., Lai, H.-T., Hsiao, C.-D., 2020. Multiple Screening of Pesticides Toxicity in Zebrafish and Daphnia Based on Locomotor Activity Alterations. Biomolecules 10, 1224. https://doi.org/10.3390/biom10091224

Jia, M., Teng, M., Tian, S., Yan, J., Meng, Z., Yan, S., Li, R., Zhou, Z., Zhu, W., 2020. Developmental toxicity and neurotoxicity of penconazole enantiomers exposure on zebrafish (Danio rerio). Environ. Pollut. Barking Essex 1987 267, 115450. https://doi.org/10.1016/j.envpol.2020.115450

Jiang, J., Chen, L., Wu, S., Lv, L., Liu, X., Wang, Q., Zhao, X., 2020. Effects of difenoconazole on hepatotoxicity, lipid metabolism and gut microbiota in zebrafish (Danio rerio). Environ. Pollut. Barking Essex 1987 265, 114844. https://doi.org/10.1016/j.envpol.2020.114844

Jin, Y., Zhu, Z., Wang, Y., Yang, E., Feng, X., Fu, Z., 2016. The fungicide imazalil induces developmental abnormalities and alters locomotor activity during early developmental stages in zebrafish. Chemosphere 153, 455–461. https://doi.org/10.1016/j.chemosphere.2016.03.085

Knapp, G., Hartung, J., 2003. Improved tests for a random effects meta-regression with a single covariate. Stat. Med. 22, 2693–2710. https://doi.org/10.1002/sim.1482

Kumar, N., Willis, A., Satbhai, K., Ramalingam, L., Schmitt, C., Moustaid-Moussa, N., Crago, J., 2020. Developmental toxicity in embryo-larval zebrafish (Danio rerio) exposed to strobilurin fungicides (azoxystrobin and pyraclostrobin). Chemosphere 241, 124980. https://doi.org/10.1016/j.chemosphere.2019.124980

Landis, S.C., Amara, S.G., Asadullah, K., Austin, C.P., Blumenstein, R., Bradley, E.W., Crystal, R.G., Darnell, R.B., Ferrante, R.J., Fillit, H., Finkelstein, R., Fisher, M., Gendelman, H.E., Golub, R.M., Goudreau, J.L., Gross, R.A., Gubitz, A.K., Hesterlee, S.E., Howells, D.W., Huguenard, J., Kelner, K., Koroshetz, W., Krainc, D., Lazic, S.E., Levine, M.S., Macleod, M.R., McCall, J.M., Moxley, R.T., Narasimhan, K., Noble, L.J., Perrin, S., Porter, J.D., Steward, O., Unger, E., Utz, U., Silberberg, S.D., 2012. A call for transparent reporting to optimize the predictive value of preclinical research. Nature 490, 187–191. https://doi.org/10.1038/nature11556

Leandro, L.P., Siqueira de Mello, R., da Costa-Silva, D.G., Medina Nunes, M.E., Rubin Lopes, A., Kemmerich Martins, I., Posser, T., Franco, J.L., 2021. Behavioral changes occur earlier than redox alterations in developing zebrafish exposed to Mancozeb. Environ. Pollut. 268, 115783. https://doi.org/10.1016/j.envpol.2020.115783

Li, H., Yu, S., Cao, F., Wang, C., Zheng, M., Li, X., Qiu, L., 2018. Developmental toxicity and potential mechanisms of pyraoxystrobin to zebrafish (Danio rerio). Ecotoxicol. Environ. Saf. 151, 1–9. https://doi.org/10.1016/j.ecoenv.2017.12.061

Li, H., Zhao, F., Cao, F., Teng, M., Yang, Y., Qiu, L., 2019. Mitochondrial dysfunction-based cardiotoxicity and neurotoxicity induced by pyraclostrobin in zebrafish larvae. Environ. Pollut. 251, 203–211. https://doi.org/10.1016/j.envpol.2019.04.122

Li, X.Y., Qin, Y.J., Wang, Y., Huang, T., Zhao, Y.H., Wang, X.H., Martyniuk, C.J., Yan, B., 2021. Relative comparison of strobilurin fungicides at environmental levels: Focus on mitochondrial function and larval activity in early staged zebrafish (Danio rerio). Toxicology 452, 152706. https://doi.org/10.1016/j.tox.2021.152706

Lin, H., Lin, F., Yuan, J., Cui, F., Chen, J., 2021. Toxic effects and potential mechanisms of Fluxapyroxad to zebrafish (Danio rerio) embryos. Sci. Total Environ. 769, 144519. https://doi.org/10.1016/j.scitotenv.2020.144519

Liu, X., Zhang, R., Jin, Y., 2020. Differential responses of larval zebrafish to the fungicide propamocarb: Endpoints at development, locomotor behavior and oxidative stress. Sci. Total Environ. 731, 139136. https://doi.org/10.1016/j.scitotenv.2020.139136

Lopes, A.R., Moraes, J.S., Martins, C. de M.G., 2022. Effects of the herbicide glyphosate on fish from embryos to adults: a review addressing behavior patterns and mechanisms behind them. Aquat. Toxicol. Amst. Neth. 251, 106281. https://doi.org/10.1016/j.aquatox.2022.106281

Lulla, A., Barnhill, L., Bitan, G., Ivanova, M.I., Nguyen, B., O’Donnell, K.C., Stahl, M.C., Yamashiro, C., Klärner F.-G., Schrader, T., Sagasti, A., Bronstein, J.M., 2016. Neurotoxicity of the Parkinson Disease-Associated Pesticide Ziram Is Synuclein-Dependent in Zebrafish Embryos. Environ. Health Perspect. 124, 1766–1775. https://doi.org/10.1289/EHP141

Lushchak, V.I., Matviishyn, T.M., Husak, V.V., Storey, J.M., Storey, K.B., 2018. Pesticide toxicity: a mechanistic approach. EXCLI J. 17, 1101–1136. https://doi.org/10.17179/excli2018-1710

Maggi, F., Tang, F.H.M., la Cecilia, D., McBratney, A., 2019. PEST-CHEMGRIDS, global gridded maps of the top 20 crop-specific pesticide application rates from 2015 to 2025. Sci. Data 6, 170. https://doi.org/10.1038/s41597-019-0169-4

McGuinness, L.A., Higgins, J.P.T., 2021. Risk-of-bias VISualization (robvis): An R package and Shiny web app for visualizing risk-of-bias assessments. Res. Synth. Methods 12, 55–61. https://doi.org/10.1002/jrsm.1411

McMahon, T.A., Halstead, N.T., Johnson, S., Raffel, T.R., Romansic, J.M., Crumrine, P.W., Rohr, J.R., 2012. Fungicide-induced declines of freshwater biodiversity modify ecosystem functions and services. Ecol. Lett. 15, 714–722. https://doi.org/10.1111/j.1461-0248.2012.01790.x

Meng, Y., Zhong, K., Xiao, J., Huang, Y., Wei, Y., Tang, L., Chen, S., Wu, J., Ma, J., Cao, Z., Liao, X., Lu, H., 2020. Exposure to pyrimethanil induces developmental toxicity and cardiotoxicity in zebrafish. Chemosphere 255, 126889. https://doi.org/10.1016/j.chemosphere.2020.126889

Miller, R.G., 1974. The Jackknife--A Review. Biometrika 61, 1–15. https://doi.org/10.2307/2334280

Mu, X., Chai, T., Wang, K., Zhu, L., Huang, Y., Shen, G., Li, Y., Li, X., Wang, C., 2016. The developmental effect of difenoconazole on zebrafish embryos: A mechanism research. Environ. Pollut. Barking Essex 1987 212, 18–26. https://doi.org/10.1016/j.envpol.2016.01.035

Mu, X., Pang, S., Sun, X., Gao, J., Chen, J., Chen, X., Li, X., Wang, C., 2013. Evaluation of acute and developmental effects of difenoconazole via multiple stage zebrafish assays. Environ. Pollut. Barking Essex 1987 175, 147–157. https://doi.org/10.1016/j.envpol.2012.12.029

OECD, 2019. Test No. 203: Fish, Acute Toxicity Test. Organisation for Economic Co-operation and Development, Paris.

OECD, 2013. Test No. 236: Fish Embryo Acute Toxicity (FET) Test. Organisation for Economic Co-operation and Development, Paris.

Ouzzani, M., Hammady, H., Fedorowicz, Z., Elmagarmid, A., 2016. Rayyan— a web and mobile app for systematic reviews. Syst. Rev. 5, 210. https://doi.org/10.1186/s13643-016-0384-4

Page, M.J., McKenzie, J.E., Bossuyt, P.M., Boutron, I., Hoffmann, T.C., Mulrow, C.D., Shamseer, L., Tetzlaff, J.M., Akl, E.A., Brennan, S.E., Chou, R., Glanville, J., Grimshaw, J.M., Hróbjartsson, A., Lalu, M.M., Li, T., Loder, E.W., Mayo-Wilson, E., McDonald, S., McGuinness, L.A., Stewart, L.A., Thomas, J., Tricco, A.C., Welch, V.A., Whiting, P., Moher, D., 2021. The PRISMA 2020 statement: an updated guideline for reporting systematic reviews. BMJ 372, n71. https://doi.org/10.1136/bmj.n71

Pang, S., Guo, M., Zhang, X., Yu, L., Zhang, Z., Huang, L., Gao, J., Li, X., 2020. Myclobutanil developmental toxicity, bioconcentration and sex specific response in cholesterol in zebrafish (Denio rerio). Chemosphere 242, 125209. https://doi.org/10.1016/j.chemosphere.2019.125209

Paredes-Zúñiga, S., Ormeño, F., Allende, M.L., 2021. Triadimefon triggers circling behavior and conditioned place preference/aversion in zebrafish in a dose dependent manner. Neurotoxicol. Teratol. 86, 106979. https://doi.org/10.1016/j.ntt.2021.106979

Paredes-Zúñiga, S., Trost, N., De la Paz, J.F., Alcayaga, J., Allende, M.L., 2019. Behavioral effects of triadimefon in zebrafish are associated with alterations of the dopaminergic and serotonergic pathways. Prog. Neuropsychopharmacol. Biol. Psychiatry 92, 118–126. https://doi.org/10.1016/j.pnpbp.2018.12.012

Perez-Rodriguez, V., Souders, C.L., Tischuk, C., Martyniuk, C.J., 2019. Tebuconazole reduces basal oxidative respiration and promotes anxiolytic responses and hypoactivity in early-staged zebrafish (Danio rerio). Comp. Biochem. Physiol. Part C Toxicol. Pharmacol. 217, 87–97. https://doi.org/10.1016/j.cbpc.2018.11.017

Pignati, W.A., Lima, F.A.N. de S.E., Lara, S.S. de, Correa, M.L.M., Barbosa, J.R., Leão, L.H. da C., Pignatti, M.G., 2017. Spatial distribution of pesticide use in Brazil: a strategy for Health Surveillance. Cienc. Saude Coletiva 22, 3281–3293. https://doi.org/10.1590/1413-812320172210.17742017

Pimentel, D., 1995. Amounts of pesticides reaching target pests: Environmental impacts and ethics. J. Agric. Environ. Ethics 8, 17–29. https://doi.org/10.1007/BF02286399

Pompermaier, A., Varela, A.C.C., Fortuna, M., Mendonça-Soares, S., Koakoski, G., Aguirre, R., Oliveira, T.A., Sordi, E., Moterle, D.F., Pohl, A.R., Rech, V.C., Bortoluzzi, E.C., Barcellos, L.J.G., 2021. Water and suspended sediment runoff from vineyard watersheds affecting the behavior and physiology of zebrafish. Sci. Total Environ. 757, 143794. https://doi.org/10.1016/j.scitotenv.2020.143794

Qian, L., Cui, F., Yang, Y., Liu, Y., Qi, S., Wang, C., 2018. Mechanisms of developmental toxicity in zebrafish embryos (Danio rerio) induced by boscalid. Sci. Total Environ. 634, 478–487. https://doi.org/10.1016/j.scitotenv.2018.04.012

Qian, L., Qi, S., Cao, F., Zhang, J., Li, C., Song, M., Wang, C., 2019. Effects of penthiopyrad on the development and behaviour of zebrafish in early-life stages. Chemosphere 214, 184–194. https://doi.org/10.1016/j.chemosphere.2018.09.117

Qian, L., Qi, S., Wang, Z., Magnuson, J.T., Volz, D.C., Schlenk, D., Jiang, J., Wang, C., 2021. Environmentally relevant concentrations of boscalid exposure affects the neurobehavioral response of zebrafish by disrupting visual and nervous systems. J. Hazard. Mater. 404, 124083. https://doi.org/10.1016/j.jhazmat.2020.124083

Reis, C.G., Herrmann, A.P., Piato, A., Bastos, L.M., Zanona, Q.K., Chitolina, R., Becker, S.Z., 2021. Neurobehavioral effects of fungicides in zebrafish: protocol. https://doi.org/10.17605/OSF.IO/F2D38

Richardson, J.R., Fitsanakis, V., Westerink, R.H.S., Kanthasamy, A.G., 2019. Neurotoxicity of pesticides. Acta Neuropathol. (Berl.) 138, 343–362. https://doi.org/10.1007/s00401-019-02033-9

Richardson, M., Garner, P., Donegan, S., 2019. Interpretation of subgroup analyses in systematic reviews: A tutorial. Clin. Epidemiol. Glob. Health 7, 192–198. https://doi.org/10.1016/j.cegh.2018.05.005

Samsa, G., Samsa, L., 2019. A Guide to Reproducibility in Preclinical Research. Acad. Med. J. Assoc. Am. Med. Coll. 94, 47–52. https://doi.org/10.1097/ACM.0000000000002351

Santana, M.S., Sandrini-Neto, L., Di Domenico, M., Prodocimo, M.M., 2021. Pesticide effects on fish cholinesterase variability and mean activity: A meta-analytic review. Sci. Total Environ. 757, 143829. https://doi.org/10.1016/j.scitotenv.2020.143829

Sert, N.P. du, Hurst, V., Ahluwalia, A., Alam, S., Avey, M.T., Baker, M., Browne, W.J., Clark, A., Cuthill, I.C., Dirnagl, U., Emerson, M., Garner, P., Holgate, S.T., Howells, D.W., Karp, N.A., Lazic, S.E., Lidster, K., MacCallum, C.J., Macleod, M., Pearl, E.J., Petersen, O.H., Rawle, F., Reynolds, P., Rooney, K., Sena, E.S., Silberberg, S.D., Steckler, T., Würbel, H., 2020. The ARRIVE guidelines 2.0: Updated guidelines for reporting animal research. PLOS Biol. 18, e3000410. https://doi.org/10.1371/journal.pbio.3000410

Sharma, A., Kumar, V., Shahzad, B., Tanveer, M., Sidhu, G.P.S., Handa, N., Kohli, S.K., Yadav, P., Bali, A.S., Parihar, R.D., Dar, O.I., Singh, K., Jasrotia, S., Bakshi, P., Ramakrishnan, M., Kumar, S., Bhardwaj, R., Thukral, A.K., 2019. Worldwide pesticide usage and its impacts on ecosystem. SN Appl. Sci. 1, 1446. https://doi.org/10.1007/s42452-019-1485-1

Shen, C., Zhou, Y., Tang, C., He, C., Zuo, Z., 2020. Developmental exposure to mepanipyrim induces locomotor hyperactivity in zebrafish (Danio rerio) larvae. Chemosphere 256, 127106. https://doi.org/10.1016/j.chemosphere.2020.127106

Souders, C.L., Perez-Rodriguez, V., El Ahmadie, N., Zhang, X., Tischuk, C., Martyniuk, C.J., 2020. Investigation into the sub-lethal effects of the triazole fungicide triticonazole in zebrafish (Danio rerio) embryos/larvae. Environ. Toxicol. 35, 254–267. https://doi.org/10.1002/tox.22862

Souders, C.L., Xavier, P., Perez-Rodriguez, V., Ector, N., Zhang, J.-L., Martyniuk, C.J., 2019. Sub-lethal effects of the triazole fungicide propiconazole on zebrafish (Danio rerio) development, oxidative respiration, and larval locomotor activity. Neurotoxicol. Teratol. 74, 106809. https://doi.org/10.1016/j.ntt.2019.106809

Tang, C., Shen, C., Zhu, K., Zhou, Y., Chuang, Y.-J., He, C., Zuo, Z., 2020. Exposure to the AhR agonist cyprodinil impacts the cardiac development and function of zebrafish larvae. Ecotoxicol. Environ. Saf. 201, 110808. https://doi.org/10.1016/j.ecoenv.2020.110808

Tang, C., Zhu, Y., Laziyan, Y., Yang, C., He, C., Zuo, Z., 2021. Long-term exposure to cyprodinil causes abnormal zebrafish aggressive and antipredator behavior through the hypothalamic–pituitary–interrenal axis. Aquat. Toxicol. 241, 106002. https://doi.org/10.1016/j.aquatox.2021.106002

Teng, M., Wang, Chen, Song, M., Chen, X., Zhang, J., Wang, Chengju, 2020. Chronic exposure of zebrafish (Danio rerio) to flutolanil leads to endocrine disruption and reproductive disorders. Environ. Res. 184, 109310. https://doi.org/10.1016/j.envres.2020.109310

Teng, M., Zhao, F., Zhou, Y., Yan, S., Tian, S., Yan, J., Meng, Z., Bi, S., Wang, C., 2019. Effect of Propiconazole on the Lipid Metabolism of Zebrafish Embryos (Danio rerio). J. Agric. Food Chem. 67, 4623–4631. https://doi.org/10.1021/acs.jafc.9b00449

Teng, M., Zhu, W., Wang, D., Qi, S., Wang, Y., Yan, J., Dong, K., Zheng, M., Wang, C., 2018a. Metabolomics and transcriptomics reveal the toxicity of difenoconazole to the early life stages of zebrafish (Danio rerio). Aquat. Toxicol. Amst. Neth. 194, 112–120. https://doi.org/10.1016/j.aquatox.2017.11.009

Teng, M., Qi, S., Zhu, W., Wang, Y., Wang, D., Dong, K., Wang, C., 2018b. Effects of the bioconcentration and parental transfer of environmentally relevant concentrations of difenoconazole on endocrine disruption in zebrafish (Danio rerio). Environ. Pollut. 233, 208–217. https://doi.org/10.1016/j.envpol.2017.10.063

Tian, S., Teng, M., Meng, Z., Yan, S., Jia, M., Li, R., Liu, L., Yan, J., Zhou, Z., Zhu, W., 2019. Toxicity effects in zebrafish embryos (Danio rerio) induced by prothioconazole. Environ. Pollut. 255, 113269. https://doi.org/10.1016/j.envpol.2019.113269

Tudi, M., Daniel Ruan, H., Wang, L., Lyu, J., Sadler, R., Connell, D., Chu, C., Phung, D.T., 2021. Agriculture Development, Pesticide Application and Its Impact on the Environment. Int. J. Environ. Res. Public. Health 18, 1112. https://doi.org/10.3390/ijerph18031112

Valadas, J., Mocelin, R., Sachett, A., Marcon, M., Zanette, R.A., Dallegrave, E., Herrmann, A.P., Piato, A., 2019. Propiconazole induces abnormal behavior and oxidative stress in zebrafish. Environ. Sci. Pollut. Res. 26, 27808–27815. https://doi.org/10.1007/s11356-019-05977-3

van Eck, N.J., Waltman, L., 2010. Software survey: VOSviewer, a computer program for bibliometric mapping. Scientometrics 84, 523–538. https://doi.org/10.1007/s11192-009-0146-3

van Eck, N.J., Waltman, L., 2007. VOS: A New Method for Visualizing Similarities Between Objects, in: Decker, R., Lenz, H.-J. (Eds.), Advances in Data Analysis, Studies in Classification, Data Analysis, and Knowledge Organization. Springer, Berlin, Heidelberg, pp. 299–306. https://doi.org/10.1007/978-3-540-70981-7_34

Vasamsetti, B.M.K., Kim, N.-S., Chon, K., Park, H.-H., 2020. Teratogenic and developmental toxic effects of etridiazole on zebrafish (Danio rerio) embryos. Appl. Biol. Chem. 63, 80. https://doi.org/10.1186/s13765-020-00566-2

Veroniki, A.A., Jackson, D., Viechtbauer, W., Bender, R., Bowden, J., Knapp, G., Kuss, O., Higgins, J.P.T., Langan, D., Salanti, G., 2016. Methods to estimate the between-study variance and its uncertainty in meta-analysis. Res. Synth. Methods 7, 55–79. https://doi.org/10.1002/jrsm.1164

Vesterinen, H.M., Sena, E.S., Egan, K.J., Hirst, T.C., Churolov, L., Currie, G.L., Antonic, A., Howells, D.W., Macleod, M.R., 2014. Meta-analysis of data from animal studies: A practical guide. J. Neurosci. Methods 221, 92–102. https://doi.org/10.1016/j.jneumeth.2013.09.010

Viechtbauer, W., 2005. Bias and Efficiency of Meta-Analytic Variance Estimators in the Random-Effects Model. J. Educ. Behav. Stat. 30, 261–293. https://doi.org/10.3102/10769986030003261

Viechtbauer, W., López-López, J.A., Sánchez-Meca, J., Marín-Martínez, F., 2015. A comparison of procedures to test for moderators in mixed-effects meta-regression models. Psychol. Methods 20, 360–374. https://doi.org/10.1037/met0000023

Wang, H., Meng, Z., Liu, F., Zhou, L., Su, M., Meng, Y., Zhang, S., Liao, X., Cao, Z., Lu, H., 2020. Characterization of boscalid-induced oxidative stress and neurodevelopmental toxicity in zebrafish embryos. Chemosphere 238, 124753. https://doi.org/10.1016/j.chemosphere.2019.124753

Wang, H., Zhou, L., Liao, X., Meng, Z., Xiao, J., Li, F., Zhang, S., Cao, Z., Lu, H., 2019. Toxic effects of oxine-copper on development and behavior in the embryo-larval stages of zebrafish. Aquat. Toxicol. Amst. Neth. 210, 242–250. https://doi.org/10.1016/j.aquatox.2019.02.020

Wang, X., Li, X., Wang, Y., Qin, Y., Yan, B., Martyniuk, C.J., 2021. A comprehensive review of strobilurin fungicide toxicity in aquatic species: Emphasis on mode of action from the zebrafish model. Environ. Pollut. Barking Essex 1987 275, 116671. https://doi.org/10.1016/j.envpol.2021.116671

Wang, X.H., Zheng, S.S., Huang, T., Su, L.M., Zhao, Y.H., Souders, C.L., Martyniuk, C.J., 2018. Fluazinam impairs oxidative phosphorylation and induces hyper/hypo-activity in a dose specific manner in zebrafish larvae. Chemosphere 210, 633–644. https://doi.org/10.1016/j.chemosphere.2018.07.056

Wilkinson, L., 2011. ggplot2: Elegant Graphics for Data Analysis by WICKHAM, H. Biometrics 67, 678–679. https://doi.org/10.1111/j.1541-0420.2011.01616.x

Worp, H.B. van der, Howells, D.W., Sena, E.S., Porritt, M.J., Rewell, S., O’Collins, V., Macleod, M.R., 2010. Can Animal Models of Disease Reliably Inform Human Studies? PLOS Med. 7, e1000245. https://doi.org/10.1371/journal.pmed.1000245

Wu, A., Yu, Q., Lu, H., Lou, Z., Zhao, Y., Luo, T., Fu, Z., Jin, Y., 2021.Developmental toxicity of procymidone to larval zebrafish based on physiological and transcriptomic analysis. Comp. Biochem. Physiol. Part C Toxicol. Pharmacol. 248, 109081. https://doi.org/10.1016/j.cbpc.2021.109081

Yang, L., Huang, T., Li, R., Souders, C.L., Rheingold, S., Tischuk, C., Li, N., Zhou, B., Martyniuk, C.J., 2021. Evaluation and comparison of the mitochondrial and developmental toxicity of three strobilurins in zebrafish embryo/larvae. Environ. Pollut. Barking Essex 1987 270, 116277. https://doi.org/10.1016/j.envpol.2020.116277

Yang, Y., Chang, J., Wang, D., Ma, H., Li, Y., Zheng, Y., 2021b. Thifluzamide exposure induced neuro-endocrine disrupting effects in zebrafish (Danio rerio). Arch. Toxicol. 95, 3777–3786. https://doi.org/10.1007/s00204-021-03158-1

Yang, Y., Dong, F., Liu, X., Xu, J., Wu, X., Zheng, Y., 2019a. Flutolanil affects circadian rhythm in zebrafish (Danio rerio) by disrupting the positive regulators. Chemosphere 228, 649–655. https://doi.org/10.1016/j.chemosphere.2019.04.207

Yang, Y., Dong, F., Liu, X., Xu, J., Wu, X., Zheng, Y., 2019b. Dysregulation of circadian rhythm in zebrafish (Danio rerio) by thifluzamide: Involvement of positive and negative regulators. Chemosphere 235, 280–287. https://doi.org/10.1016/j.chemosphere.2019.06.153

Yang, Y., Qi, S., Wang, D., Wang, K., Zhu, L., Chai, T., Wang, C., 2016a. Toxic effects of thifluzamide on zebrafish (Danio rerio). J. Hazard. Mater. 307, 127–136. https://doi.org/10.1016/j.jhazmat.2015.12.055

Yang, Y., Qi, S., Chen, J., Liu, Y., Teng, M., Wang, C., 2016b. Toxic Effects of Bromothalonil and Flutolanil on Multiple Developmental Stages in Zebrafish. Bull. Environ. Contam. Toxicol. 97, 91–97. https://doi.org/10.1007/s00128-016-1833-4

Yanicostas, C., Soussi-Yanicostas, N., 2021. SDHI Fungicide Toxicity and Associated Adverse Outcome Pathways: What Can Zebrafish Tell Us? Int. J. Mol. Sci. 22, 12362. https://doi.org/10.3390/ijms222212362

Zhang, C., Zhou, T., Xu, Y., Du, Z., Li, B., Wang, Jinhua, Wang, Jun, Zhu, L., 2020. Ecotoxicology of strobilurin fungicides. Sci. Total Environ. 742, 140611. https://doi.org/10.1016/j.scitotenv.2020.140611

Zhang, X., Zhang, P., Perez-Rodriguez, V., Souders, C.L., Martyniuk, C.J., 2020. Assessing the toxicity of the benzamide fungicide zoxamide in zebrafish (Danio rerio): Towards an adverse outcome pathway for beta-tubulin inhibitors. Environ. Toxicol. Pharmacol. 78, 103405. https://doi.org/10.1016/j.etap.2020.103405

Zhou, Y., Chen, X., Teng, M., Zhang, J., Wang, C., 2019. Toxicity effects of captan on different life stages of zebrafish (Danio rerio). Environ. Toxicol. Pharmacol. 69, 80–85. https://doi.org/10.1016/j.etap.2019.04.003

Zubrod, J.P., Bundschuh, M., Arts, G., Brühl, C.A., Imfeld, G., Knäbel, A., Payraudeau, S., Rasmussen, J.J., Rohr, J., Scharmüller, A., Smalling, K., Stehle, S., Schulz, R., Schäfer, R.B., 2019. Fungicides: An Overlooked Pesticide Class? Environ. Sci. Technol. 53, 3347–3365. https://doi.org/10.1021/acs.est.8b04392

